# Mesenchymal Stromal Cell Aging Impairs the Self-Organizing Capacity of Lung Alveolar Epithelial Stem Cells

**DOI:** 10.1101/2021.03.05.434121

**Authors:** Diptiman Chanda, Mohammad Rehan, Samuel R. Smith, Kevin G. Dsouza, Yong Wang, Karen Bernard, Deepali Kurundkar, Vinayak Memula, Kojima Kyoko, James A. Mobley, Gloria Benavides, Victor Darley-Usmar, Kim Young-il, Jaroslaw W. Zmijewski, Jessy S. Deshane, Victor J. Thannickal

## Abstract

Multicellular organisms maintain structure and function of tissues/organs through emergent, self-organizing behavior. In this report, we demonstrate a critical role for lung mesenchymal stromal cell (L-MSC) aging in determining the capacity to form 3-dimentional organoids or “alveolospheres “ with type 2 alveolar epithelial cells (AEC2s). In contrast to L-MSCs from aged mice, young L-MSCs support the efficient formation of alveolospheres when co-cultured with young or aged AEC2s. Aged L-MSCs demonstrated features of cellular senescence, altered bioenergetics, and a senescence-associated secretory profile (SASP). The reactive oxygen species generating enzyme, NADPH oxidase 4 (Nox4), was highly activated in aged L-MSCs and Nox4 downregulation was sufficient to, at least partially, reverse this age-related energy deficit, while restoring the self-organizing capacity of alveolospheres. Together, these data indicate a critical role for cellular bioenergetics and redox homeostasis in an organoid model of self-organization, and supports the concept of thermodynamic entropy in aging biology.

## Introduction

Substantial progress has been made in our understanding the biology of aging, and these advances have the potential to improve both healthspan and lifespan, while alleviating the burden of age-related diseases. Self-organization in biological systems is a process by which cells reduce their internal entropy and maintain order within these dynamic, self-renewing systems (Chatterjee et al., 2017, Kiss et al., 2009). Exhaustion of stem/progenitor cells, cellular senescence, and altered intercellular communication have been proposed as aging hallmarks that increase susceptibility to age-related disorders(Schultz and Sinclair, 2016). However, the interactions between these hallmarks, and whether cellular bioenergetics associated with cellular senescence may account for age-associated stem cell dysfunction and altered cell-cell communication have not been well defined. In this study, we utilized an organoid model to study stem cell behavior and intercellular communication that may account for age-related phenotypes; through these studies, we identify cellular bioenergetics and redox imbalance as critical drivers of these inter-dependent aging hallmarks.

## Results and discussion

The mammalian lung serves an essential role in organismal metabolism, uniquely by serving as the primary organ for systemic exchange of oxygen for carbon dioxide. Regeneration and maintenance of structure-function of the lung are dependent on adult, tissue-resident stem cells that reside in unique niches along the airways (Basil et al., 2020, Hogan et al., 2014). Type 2 alveolar epithelial cells (AEC2s) serve as facultative stem/progenitor cells in adult mammalian lungs and differentiate into type 1 alveolar epithelial cells (AEC1s) in response to diverse injuries to reconstitute and reestablish the alveolar gas exchange surface (Barkauskas et al., 2013). AEC2 regenerative capacity declines with age resulting in impaired lung injury repair responses, thus increasing susceptibility to various lung diseases (Schulte et al., 2019, Watson et al., 2020). To further explore the relationship between AECs and L-MSCs, we developed an alveolosphere assay system which has been traditionally used to assess AEC2 stemness or regenerative potential (Barkauskas et al., 2013). In this assay system, L-MSCs and AEC2s are mixed together in a ratio of 100,000 to 5000, respectively, and seeded in Matrigel (**Figure 1A**); alveolospheres with a single layer of epithelial cells composed of both AEC2s [surfactant protein C (SFTPC)-positive] and AEC1s [lung type-I integral membrane glycoprotein (T1α)-positive] surrounding a hollow sphere typically form 9-12 days following co-culture (**Figure 1B, C**). L-MSCs expressing platelet-derived growth factor receptor-α (PDFGRα), with a fewer number expressing α-smooth muscle actin (α-SMA), were found primarily in cells lining the outer edges of alveolospheres (**Figure 1D, E**). AEC2s within alveolospheres stained for both the cell proliferation marker, Ki-67, and the double strand DNA damage repair marker, histone H2A.X, suggesting ongoing AEC2 turnover within these 3D organoids (**Figure 1, F-G**). This self-organizing behavior of L-MSCs and AEC2s is critically dependent on the presence both cell types, as AEC2s in the absence of AEC2s do not self-organize to form alveolospheres (**Figure 1**-**figure supplement 1A**). Together, these data support the crosstalk between L-MSC and AEC2s that permit formation of distinct alveolar organoid-like structures; this intercellular communication may be perturbed during aging accounting for age-associated loss of AEC2 regeneration/maintenance.

**Figure 1:**
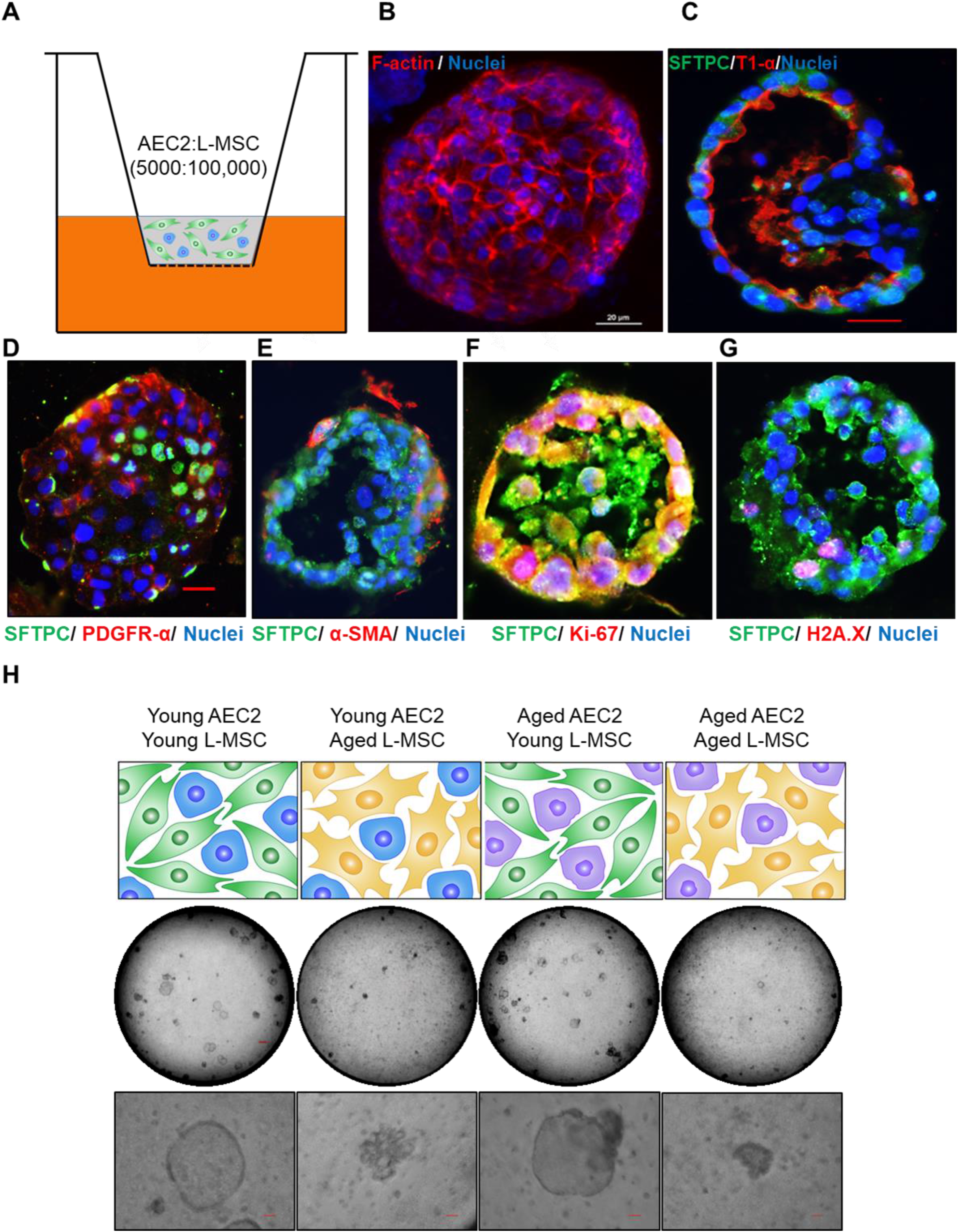
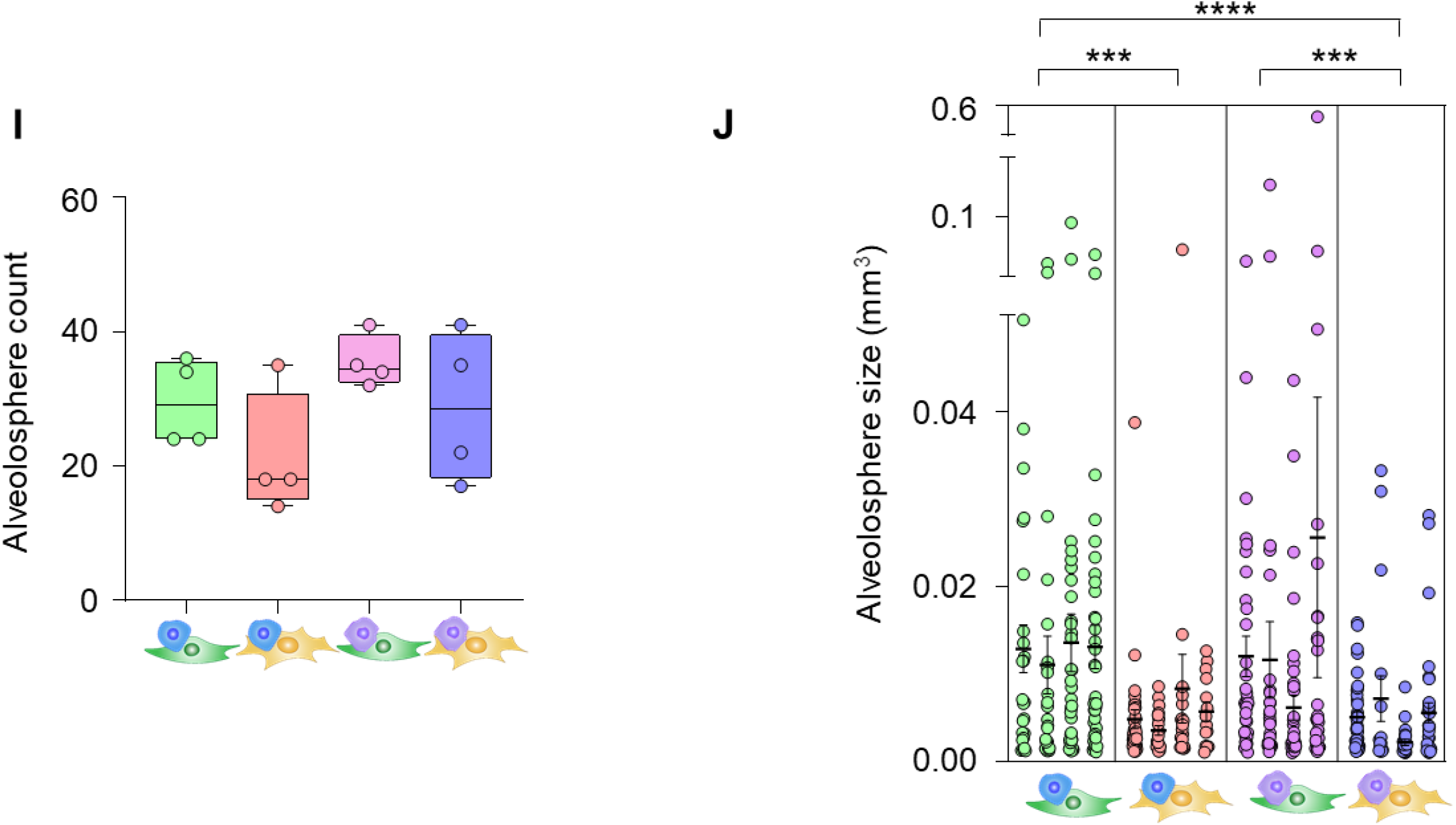
Aging lung-mesenchymal stromal cells (L-MSCs) impair self-organization of alveolar epithelial stem cells (AEC2s) and alveolosphere formation. (**A**) Alveolosphere assay. AEC2s and L-MSCs were purified from the young (3 months) mice lungs. AEC2s (5,000 cells/well) and L-MSCs (100,000 cells/well) were mixed and seeded in Matrigel: MTEC plus media (1:1) and co-cultured in cell-culture inserts in 24-well dishes as shown. Alveolospheres form within 9-12 days of co-culture. **(B)** Alveolosphere whole-mounts were immunofluorescently stained with antibody against F-actin (red) for confocal imaging. Image showing maximum intensity projection of an alveolosphere (scale bar = 20 µm). Immunofluorescence (IF) staining showing localization of AEC2s [surfactant protein-C (SFTPC), green] and AEC1s [lung type-I integral membrane glycoprotein-α (T1-α), red] within the alveolospheres. IF staining showing localization of platelet-derived growth factor receptor-α (PDGFR-α, red) expressing alveolar L-MSCs within the alveolospheres. (**E**) Alpha-smooth muscle actin (α-SMA, red) IF staining showing presence of myofibroblasts in the alveolosphere. (**F-G**) Cell proliferation and DNA repair/apoptosis within the alveolospheres were determined by Ki-67 (**F**; red) and Histone H2A.X (**G**; red) IF staining respectively. Nuclei were stained with Hoechst 33342 (Blue; scale bars = 20 µm). (**H**) L-MSCs and AEC2s from young (3 months) and aged mice (24 months) were co-cultured in varied combinations as shown (upper panel); alveolospheres were imaged by brightfield microscopy. Images at low and high magnifications are shown (middle panel, scale bar = 300 µm; lower panel, scale bar = 20 µm). (**I**) Alveolospheres in each well were counted (n = 4); box and whiskers plot showing median alveolosphere count in each group (*p > 0*.*05;* one-way ANOVA; Tukey ‘s multiple comparison test). (**J**) Alveolosphere sizes (volumes) were determined for each of the co-culture groups using Image J 1.47v software. Nested scatterplot showing Mean ± SEM of all the alveolospheres counted in each well for each group (n = 4; \*\*\**p < 0*.*001*, young AEC2s:young L-MSCs *vs*. young AEC2s:aged L-MSCs; aged AEC2s:young L-MSCs *vs*. aged AEC2s:aged L-MSCs; \*\*\*\**p < 0*.*0001*, young AEC2s:young L-MSCs *vs*. aged AEC2s: aged L-MSCs; one-way ANOVA followed by Tukey ‘s multiple comparison test).

To determine the effect of age on cellular self-organization and alveolosphere formation, L-MSCs and AEC2s were isolated from the lungs of young (2-3 months) and aged (22-24 months) mice and seeded in various combinations (**Figure 1H**, upper panel). L-MSCs and AEC2s from young mice generated normal alveolospheres, whereas L-MSCs and AEC2s from the lungs of aged mice showed impaired self-organization with condensed, poorly-formed organoids. Interestingly, combining aged AEC2s with young L-MSCs resulted in relatively normal-appearing alveolospheres, while the reverse combination (young AEC2s and old L-MSCs) did not (**Figure 1H**, middle and lower panel; **figure supplement 1B**). Quantitative analyses revealed that, while the number of alveolospheres formed were not significantly different between groups (**Figure 1I**), the size of alveolospheres was critically dependent on the age of L-MSCs, and not that of AEC2s (**Figure 1J**). These findings indicate that secreted factor(s) from young L-MSCs are capable of supporting the self-organizing behavior of alveolospheres, while aged L-MSCs do not.

Cellular senescence accumulates in tissues with advancing age (Krishnamurthy et al., 2004), and has been proposed as a key driver of aging and aging-related disease phenotypes (Baker et al., 2016, Kennedy et al., 2014). Based on our observation that aged L-MSCs were incapable of supporting alveolosphere formation, we explored whether the emergence of cellular senescence may account for this finding. First, we confirmed the senescent features of L-MSCs from aged mice in monolayer 2D cell culture (**Figure 2A-figure supplement 2A**) and in 3D alveolospheres (**Figure 2B**), by staining for β-galactosidase (β-gal) (Dimri et al., 1995) and lipofuscin (Georgakopoulou et al., 2013), respectively. To characterize secreted factors that may account for the age-associated dysregulation of cell-cell communication, we first measured cytokines/growth factors secreted from both young and aged L-MSCs using antibody arrays (**Figure 2C**). A number of cytokines that have been associated with the senescence-associated secretory phenotype (SASP) were found to be released at higher levels by aged L-MSCs, including IL-6, CCL12/MCP-5, CCL11/Eotaxin, and WISP-1/CCN4; in contrast, IGFBP1 was the only secreted protein that was statistically more elevated in young L-MSCs in this cytokine array (**Figure 2D, E**).

**Figure 2:**
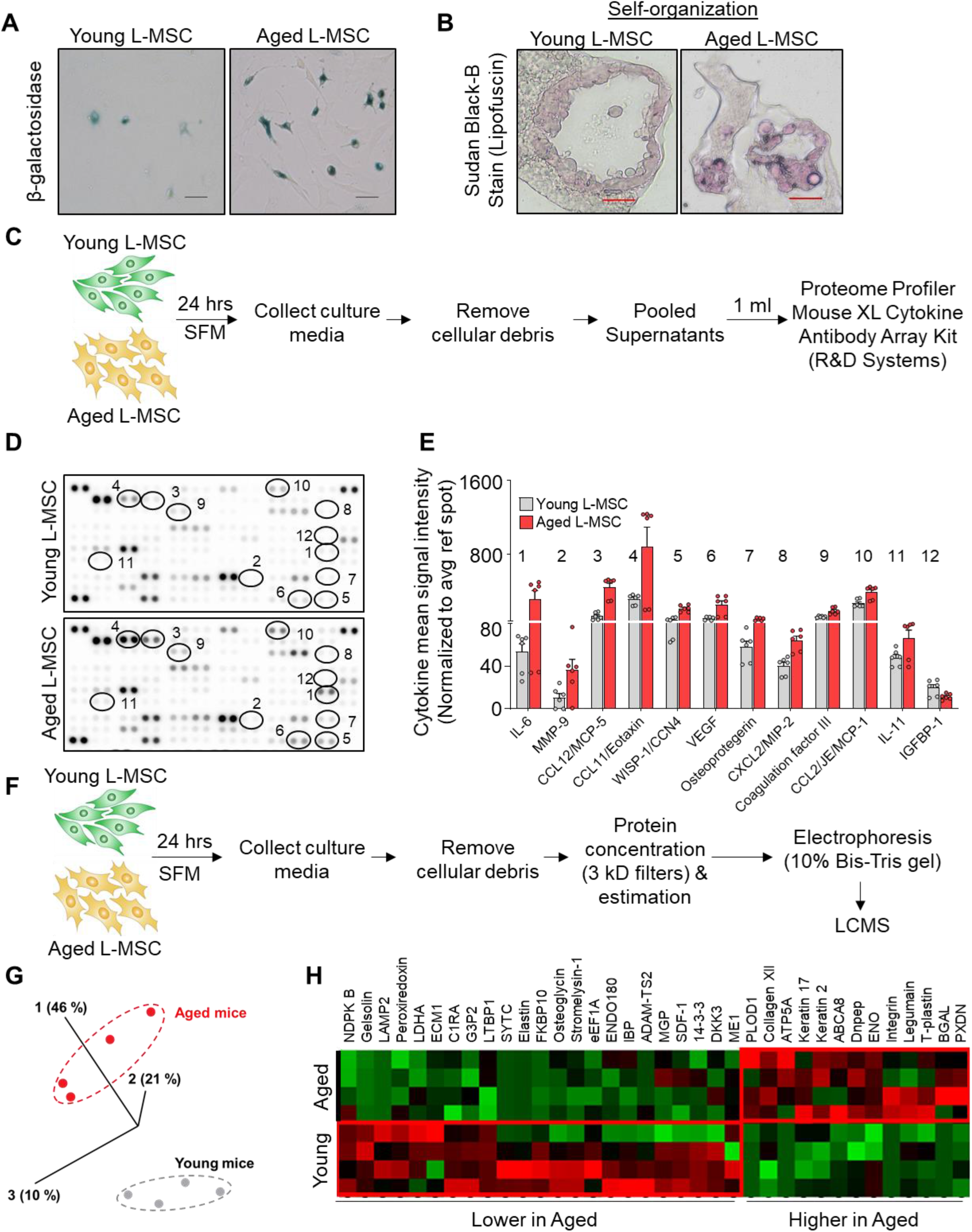
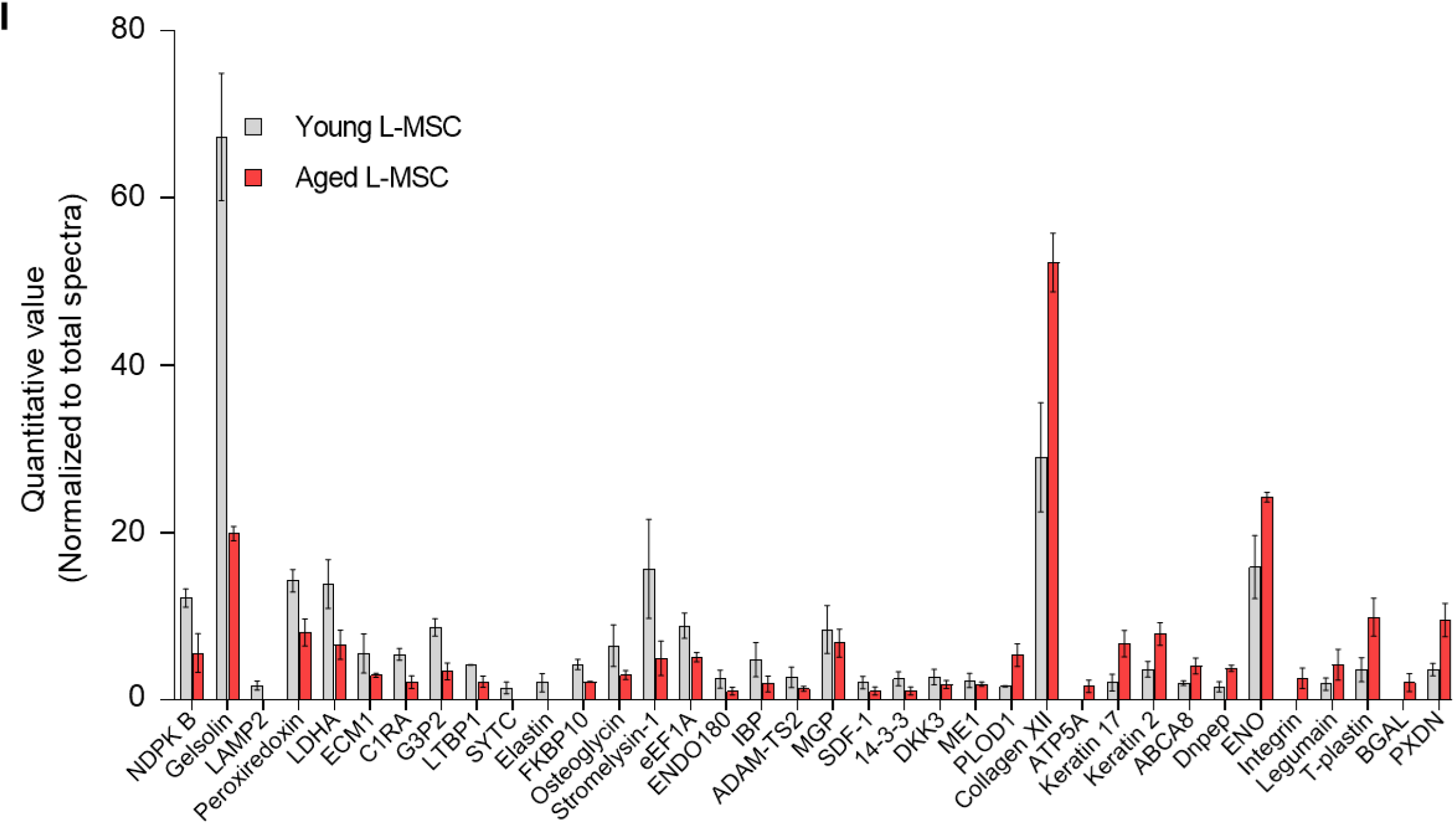
Aged L-MSCs show redox imbalance and acquire senescence-associated secretory phenotype. (**A**) β-galactosidase activity assay. Young and aged L-MSCs were plated in 6-well dishes at a density of 150,000 L-MSCs/well. L-MSCs were allowed to attach overnight and stained for β-galactosidase activity (scale bar = 50µm). (**B**) Lipofuscin granules were detected in the alveolosphere paraffin sections by Sudan Black B staining. Sections were also stained with nuclear fast red for contrast (scale bar = 20µm). (**C**) Cytokine array. 150,000 young and aged L-MSCs were plated in each well of a 6-well culture dish and grown overnight. Cells were washed with PBS and cultured for 24 hrs in serum-free media (SFM; 1.5 ml/well). The culture media were pooled for each cell type and centrifuged to remove any cell and debris. 1 ml of supernatant was applied to antibody array, dotted with antibodies against 111 mouse cytokines and growth factors in duplicates. (**D**) Antibody array showing comparative expression of cytokines and growth factors in the culture media obtained from young and aged L-MSCs. Cytokines and growth factors showing significant difference are numbered and circled. (**E**) Signal intensity was determined for each dot using Image Quant array analysis software; mean signal intensity was calculated for each cytokine, and plotted. Twelve proteins with statistically significant difference (n = 6; *p < 0*.*05*; unpaired T-test) between the young and aged L-MSCs are shown. Data presented here include pooled data from 3 independent experiments. Proteomics/mass-spectrometry analysis. Cell culture media were collected from young and aged L-MSCs (10^6^ cells) after 24 hrs of growth in SFM in 10 cm dishes, and subjected to proteomics analysis by liquid chromatography-mass spectrometry (LCMS). (**G**) A three-dimensional principal component analysis (PCA) plot showing replicated samples (young and aged) are relatively similar in their protein expression profiles and grouped together. (**H**) Heat map showing comparative expression of highly secreted proteins in the culture media between young and aged L-MSCs (n = 4). (**I**) The top 36 proteins showing statistically significant difference (n = 4; *p < 0*.*05*; unpaired T-test) between the young and aged L-MSC secretome are plotted.

In addition to pre-defined cytokine array analyses, we employed an unbiased approach to identification of secreted proteins from young and aged L-MSCs by mass spectrometry-based discovery proteomics (**Figure 2F**). After adjustments for false discovery rates, 503 high confidence proteins were identified; of these, 235 proteins were found to have non-zero quantifiable values in at least 3 of 4 experimental repeats per group for subsequent statistical analysis. The 36 proteins that passed both a single pairwise statistical test (*p < 0*.*05*) in addition to a fold change of ± 1.5 (**Figure 2**-**Table supplement 1**) were subjected to principal component analysis (PCA) (**Figure 2G**) and heat map analysis (**Figure 2H**; quantitation of heat map proteins, **Figure 2I)**. Gene ontology localization analyses revealed enrichment in secreted proteins, including extracellular vesicles and extracellular matrix (**Figure 2-table supplement 2**); interestingly, gene ontology processes analysis indicated the top cellular processes as negative regulation of reactive oxygen species metabolic process and extracellular matrix/structure organization (**Figure 2-table supplement 3**). Enrichment by top toxic pathologies revealed proteins associated with lung fibrosis (**Figure 2-table supplement 4**), a disease associated with aging(Thannickal et al., 2014). Network analysis revealed upregulation of pathways associated with the canonical WNT-signaling pathway and cell cycle regulation (**Figure 2-table supplement 5**). Together, these findings suggest that aged L-MSCs are characterized by cellular senescence associated with SASP factors, oxidative stress and alterations in regenerative pathways that may account for impaired alveolosphere formation.

Altered cellular metabolism have been linked to both senescence and aging (Lopez-Otin et al., 2016, Finkel, 2015, Sun et al., 2016). To compare the energy state between young and aged lung L-MSCs, rates of mitochondrial respiration and glycolysis were analyzed by Seahorse XF Analyzer. Real-time oxygen consumption rates (OCR) and extracellular acidification rates (ECAR) were significantly higher in aged L-MSCs as compared to young L-MSCs (**Figure 3A, B**). Aged L-MSCs showed significantly higher basal, maximal, ATP-linked, proton leak, and non-mitochondrial respiration when compared to young L-MSCs with no change in reserve capacity (**Figure 3, C-H**). Higher non-mitochondrial OCR in aged L-MSCs suggest potential activation of oxygen-metabolizing NADPH oxidases in aged L-MSCs. An energy map profile of the OCR and ECAR data indicated a higher basal rate of both mitochondrial respiration and glycolysis (**Figure 3I)**, suggesting that aged L-MSCs are under a basal state of higher metabolic demand. This higher energy demand under conditions of equivalent nutrient supply was associated with reduced levels of ATP/ADP under steady-state basal conditions (**Figure 3J**). Measurements of ECAR indicated a trend towards higher rates of basal, maximal, and non-glycolysis related ECAR in aged L-MSCs compared to young L-MSCs, although these did not reach statistical significance (**Figure 3-figure supplement 2b**). Together, these data indicate that senescence of L-MSCs are characterized by higher baseline metabolic demand with reduced bioenergetic efficiency.

**Figure 3:**
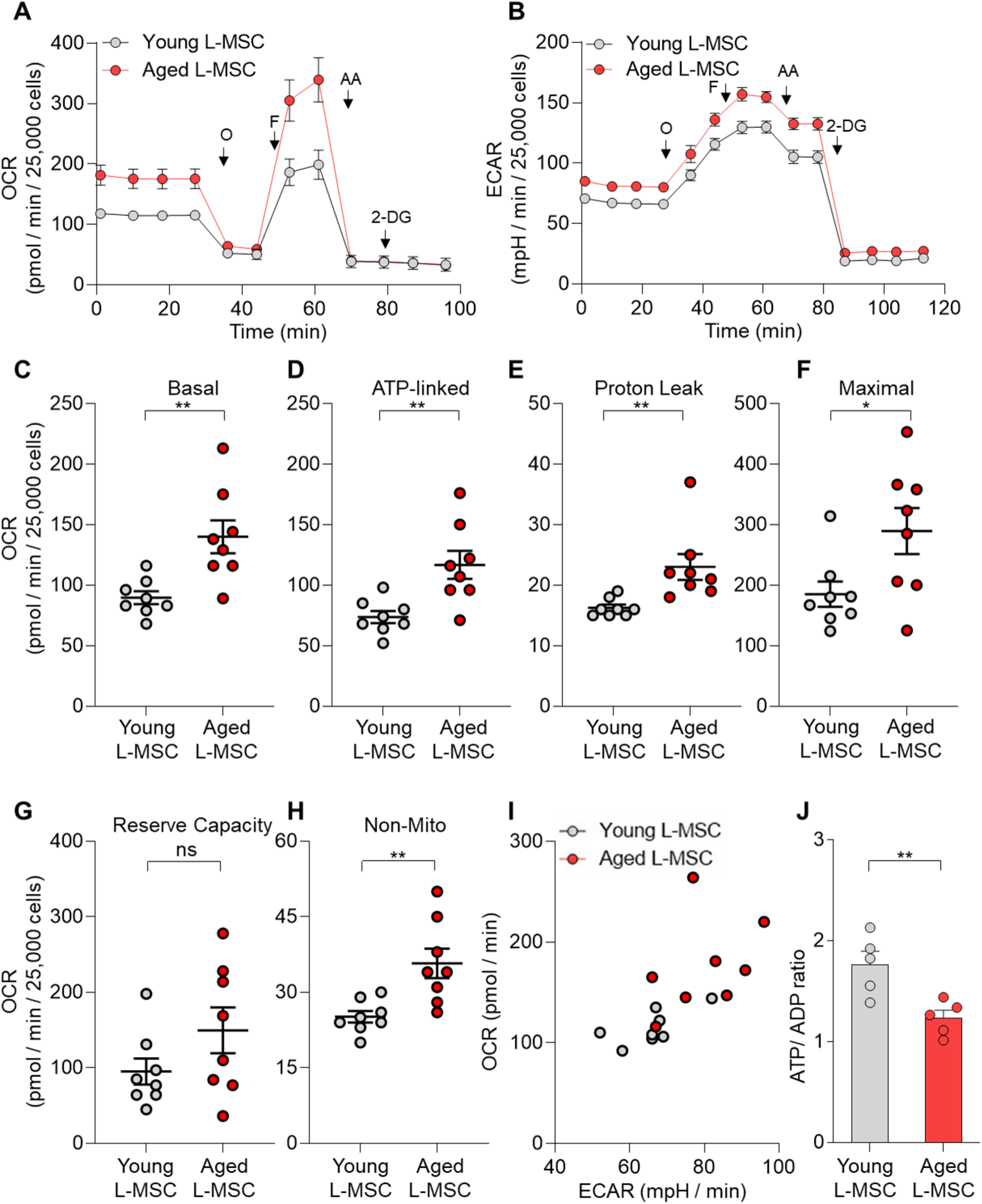
Senescent L-MSCs demonstrate altered bioenergetics. Young and aged mouse L-MSCs were grown in complete DMEM for 24 hrs. 25,000 cells were seeded in Seahorse XF-24 cell culture microplates. The cells were treated sequentially with mitochondrial inhibitors: oligomycin (Oligo), carbonyl cyanide 4-(trifluoromethoxy) phenylhydrazone (FCCP), antimycin (AA), and glycolytic inhibitor: 2-Deoxy-d - glucose (2-DG). (**A**) Real-time oxygen consumption rates (OCRs) and (**B**) real-time extracellular acidification rates (ECAR) between the young and aged L-MSCs were compared. (**C-H**) Basal, ATP-linked, proton leak, maximal, reserve capacity, and non-mitochondria related OCRs were calculated and plotted (n = 8; \**p < 0*.*05*, \*\**p < 0*.*01*; unpaired T-test; vs. Young L-MSCs). (**I**) An energy map was generated from the OCR and ECAR data (above) showing higher basal rate of both mitochondrial respiration and glycolysis in aged L-MSCs. (**J**) ATP/ADP ratio. Young and aged L-MSCs (10,000 cells) were plated in 96-well flat-bottomed dish and allowed to attach. Cells were treated for 5 min with nucleotide-releasing buffer. Relative ATP and ADP levels were measured from luminescent conversion of ATP-dependent luciferin by the luciferase enzyme. ATP/ADP ratio was calculated and plotted. Graph showing comparative ATP/ADP ratio between young and aged L-MSCs (n = 5; \*\**p < 0*.*01*; unpaired T-test; vs. young L-MSCs).

Hydrogen peroxide (H_2_O_2_) has emerged as critical regulator of redox signaling and oxidative stress (Sies, 2017). Replication-induced senescence of human lung fibroblasts results in higher rates of extracellular H_2_O_2_ release in association with increased expression of NADPH oxidase 4 (Nox4) (Sanders et al., 2015), a gene that is inducible by the pro-senescent/pro-fibrotic mediator, transforming growth factor-β1 (TGF-β1) (Hecker et al., 2009). We explored whether L-MSCs isolated from naturally aged mice are associated with higher Nox4-dependent H_2_O_2_ secretion; in comparison to young (3 months old) L-MSCs, aged (22-24 months old) L-MSCs released higher baseline levels of H_2_O_2_, an effect that was further stimulated by exogenous stimulation with TGF-β1 (**Figure 4A**). This age-dependent, pro-oxidant phenotype of L-MSCs was markedly reduced in aged mice heterozygous for Nox4 (Nox4^-/+^) (**Figure 4B, C**), implicating Nox4 as a critical mediator of both basal and TGF-β1-induced H_2_O_2_ release.

**Figure 4:**
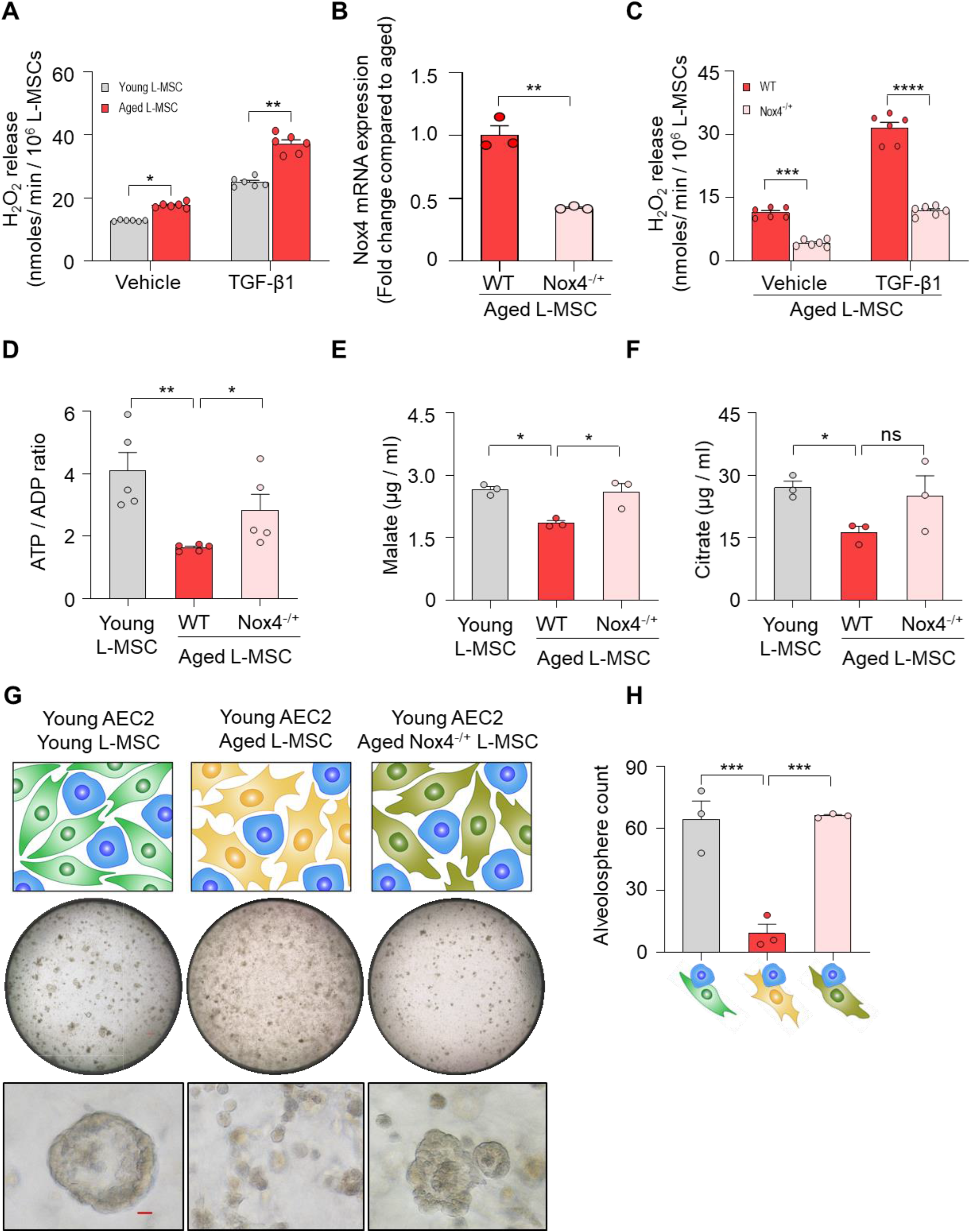
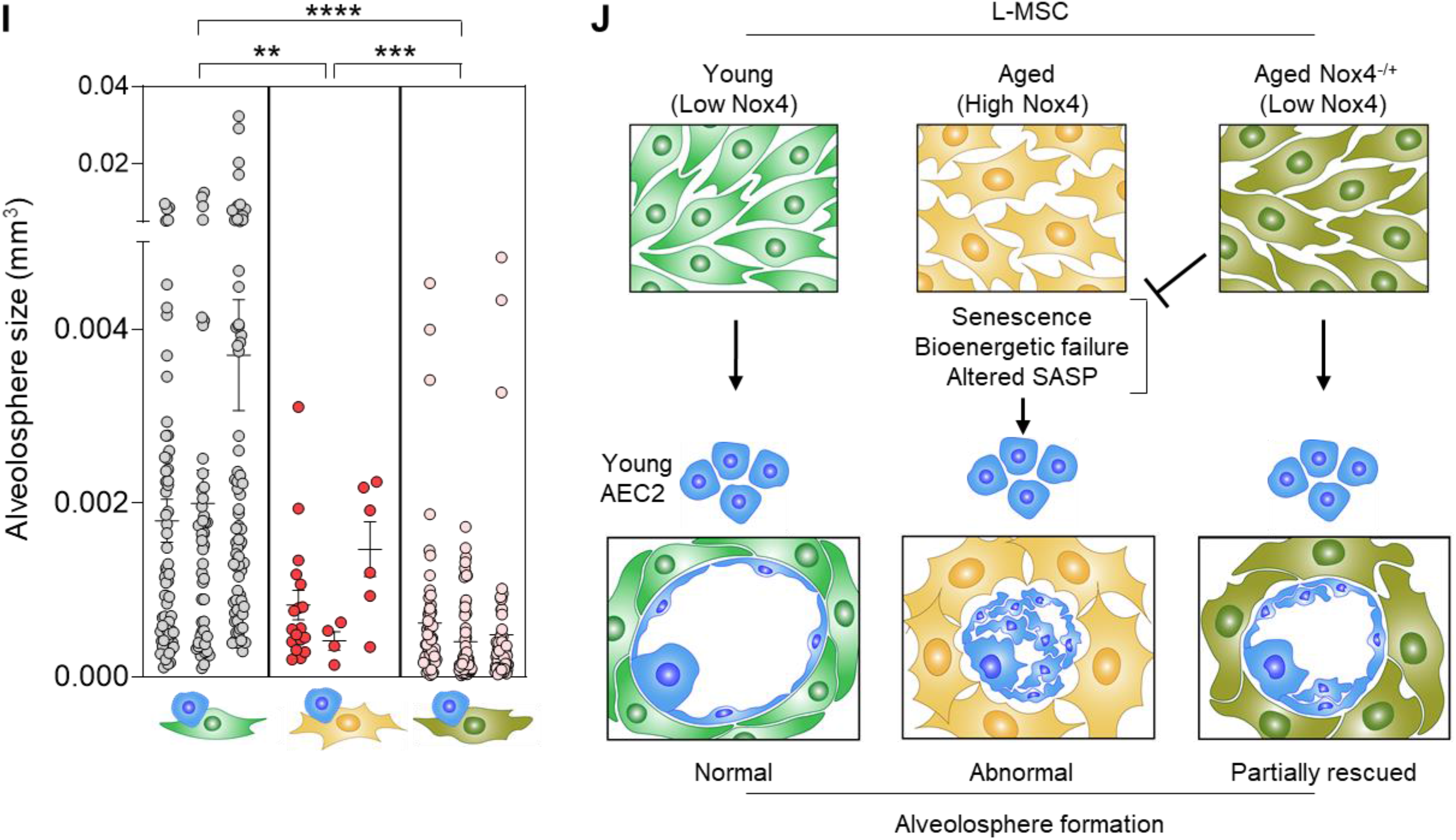
Nox4-deficiency in aged L-MSC improves bioenergetics and restores AEC2 self-organization. (**A**) Hydrogen peroxide (H2O2)-release assay. Young and aged L-MSCs were grown in very low serum (1%) containing media for 24 hrs, and treated with either transforming growth factor-β1 (TGF-β1) or vehicle for 16 hrs, and real-time H2O2 release was determined. Here, bar graph showing comparative H2O2 release between young and aged L-MSCs with or without TGF-β1 treatment (n = 6; \**p < 0*.*05*, \*\**p < 0*.*01*; unpaired T-test). (**B**) RT-PCR analysis. Total RNA was isolated from aged, and aged Nox4^-/+^ L-MSCs, and subjected to real-time PCR analysis for Nox4 mRNA expression. Data were normalized to β-actin and represented graphically as fold change compared to aged L-MSCs (n = 3; \*\**p < 0*.*01*; unpaired T-test). (**C**) Data showing comparative H2O2 release between aged, and aged Nox4^-/+^ L-MSCs at baseline and after TGF-β1 treatment (n = 6; \*\*\**p < 0*.*001*, \*\*\*\**p < 0*.*0001*; unpaired T-test). (**D**) Bar graph showing comparative ATP/ADP ratio between young, aged, and aged Nox4^-/+^ L-MSCs (n = 5; \*\**p < 0*.*01*, \**p < 0*.*05*). (**E-F**) Targeted metabolomics. Relative levels of tricarboxylic acid cycle metabolites were determined and compared between young, aged, and aged-Nox4^-/+^ L-MSCs. Concentrations of malate (**E**) and citrate (**F**) are shown here (n = 3; \**p < 0*.*05*). (**G**) Alveolosphere assay. Young, aged, and aged Nox4^-/+^ L-MSCs were co-cultured with young AEC2s in a ratio described earlier (upper panel). The alveolospheres were imaged by brightfield microscopy after 12 days of co-culture, and comparative outcomes are shown here in low (middle panel; scale bar = 300 µm) and higher magnifications (lower panel; scale bar = 20 µm). (**H**) Alveolospheres in each well were counted (Mean ± SEM; n = 3; \*\*\**p < 0*.*001*). (**I**) Alveolosphere size (volumes) were determined for each of the 3 co-culture groups. Nested scatterplot showing Mean ± SEM of all the alveolospheres counted in each well for each group (n = 3; \*\**p < 0*.*01*, \*\*\**p < 0*.*001*, \*\*\*\**p < 0*.*0001*). (**D-I**) Statistical analysis: one-way ANOVA followed by Tukey ‘s multiple comparison test. (**J**) Schematic summarizing the important findings from this study.

Recent studies implicate Nox4 in metabolic reprogramming (Bernard et al., 2017), although effects of Nox4 on age-related metabolic dysfunction are unclear. The reduced levels of ATP/ADP in aged L-MSCs were partially rescued in age-matched Nox4^-/+^ L-MSCs (**Figure 4D**), implicating a role for Nox4 in the reduced bioenergetic efficiency associated with aging. The steady-state levels of the tricarboxylic acid (TCA) cycle metabolites, malate and citrate, were reduced in wild-type aged L-MSCs; these levels recovered to levels similar to young L-MSCs in Nox4^-/+^ L-MSCs, although only malate achieved statistical significance (**Figure 4E, F**). Levels of other TCA metabolites that were not significantly altered with aging in L-MSCs were not affected by a deficiency in Nox4 (**Figure 4**-f**igure supplement 2, C-F**). These studies indicate that the Nox4 contributes to metabolic dysfunction and bioenergetic degeneration with aging in L-MSCs. Based on the observations that Nox4 mediates effects on cellular bioenergetics of L-MSCs, we examined effects of Nox4 on regulating the self-organizing potential of alveolospheres. The aberrant formation of alveolospheres when young AEC2s were cultured with aged L-MSCs were rescued when replaced with age-matched Nox4^-/+^ L-MSCs (**Figure 4G; figure supplement 3**); quantitation of both alveolospheres counts (**Figure 4H**) and alveolosphere size (**Figure 4I**) were significantly enhanced when these assays were conducted with Nox4-deficient L-MSCs. Together, our studies support a critical role for Nox4-dependent oxidative stress, senescence and bioenergetic insufficiency in restricting the capacity for cellular self-organization in a 3D organoid model of aging and stem/progenitor cell function (**Figure 4J**).

A fundamental difference between non-living matter and life forms on our planet is the ability to extract energy from the environment, primarily through oxidation of carbon-based fuels. Thus, the second law of thermodynamics which governs both living and non-living matter does not strictly apply to living organisms capable of self-renewal through the continuous and dynamic exchange of energy (from exogenous nutrients) and mass (biosynthetic and degradative processes in respiring tissues/organs). It follows then that, when this capacity for energy-dependent self-renewal is diminished, the law of entropic degeneration may well be operative in living organisms. Aging is associated with derangements in metabolism that influences, and may in fact control, many of the well-recognized hallmarks of aging, including cellular senescence (Lopez-Otin et al., 2016, Lopez-Otin et al., 2013). The inter-relatedness of metabolism with these traditional aging hallmarks have been difficult to untangle due to complexities of studies in living organisms and the relative simplicity of 2D cell culture models.

Organoids offer the opportunity to study complex living phenomena such as emergent properties and self-organization in biological systems while, at the same time, allowing for reductionist approaches that provide mechanistic understanding of these complex processes. Using a combination of methods that included a 3D organoid model, biochemical approaches, proteomics, and gene targeting, we demonstrate a critical role for the oxygen metabolizing enzyme, Nox4, in regulating cellular bioenergetics and senescence that limits stem cell function. Consistent with contemporary theories on aging, Nox4 may function as an antagonistically pleiotropic gene and contribute a number of age-related degenerative disorders, including fibrotic diseases (Thannickal, 2010, Lambeth, 2007)). Interestingly, proteomic analyses of proteins secreted by aged L-MSCs in the current study suggested alterations in cellular redox/oxidative stress (from gene ontology analysis) and lung fibrosis (as a “toxic pathology “). These findings, as well as candidate SASP factors, highlight the emerging importance of intercellular communication and stem cell exhaustion as important hallmarks of aging (Lopez-Otin et al., 2013).

Regenerative mechanisms in adult, mammalian organisms primarily rely on the activation and differentiation of tissue/organ-resident stem cells (Hogan et al., 2014). Homeostatic maintenance of these stem cells require a tightly-regulated niche that includes other supporting cell types, in particular stromal cells (Basil et al., 2020). An important finding from our studies was the observation that L-MSC aging, rather than AEC aging, accounts for the age-related inability to form alveolospheres. Thus, aging of the stem cell niche may be as important, or even more decisive, in promoting certain age-related pathologies. Therapeutic targeting of metabolic aging within the stem cell niche, specifically that of lung-resident fibroblasts/myofibroblasts, may offer a more effective and feasible strategy for age-related degenerative disorders such as tissue/organ fibrosis.

## Materials and Methods

### Key resources

All reagents, antibodies, assay kits and mice used in this study are listed in the t**able supplement 6**. All animal protocols were approved by the Institutional Animal Care and Use Committees (IACUC) at the University of Alabama at Birmingham. The mice were acclimatized in the animal facility at least for a week before experiments. Male mice were used in this study for their greater susceptibility to age-related diseases.

### Cell culture

Mouse lung L-MSCs were isolated and propagated following protocol developed in our laboratory (Vittal et al., 2005). L-MSCs were isolated by collagenase digestion of the lungs and anchorage-dependent *ex vivo* growth on plastic culture dishes in DMEM supplemented with 10% fetal bovine serum, 4 mM L-Glutamine, 4.5 g/L glucose, 100 U/ml penicillin, 100 μg/ml streptomycin, and 1.25 μg/ml amphotericin B (Fungizone), in a humified chamber at 37°C in 5% CO2, 95% air. AEC2s were isolated and purified from the young and aged mice lungs via enzymatic digestion (Dispase II), cell-specific antibody labeling, and magnetic separation following published protocols (Sinha and Lowell, 2016) and used immediately. AEC2 purity was determined by fluorescent detection of EpCam^+^/SFTPC^+^/CD45^-^/CD31^-^ cell population using flow cytometry (Sinha and Lowell, 2016, Bertoncello and McQualter, 2011). L-MSCs and AEC2s were seeded in Matrigel mixed with MTEC/plus media(You et al., 2002) (1:1) in cell culture inserts (24-well format, 0.4 µm pore size), and co-cultured in MTEC/Plus media as shown in **Figure1. Immunofluorescence staining**. Matrigel containing alveolospheres were fixed in 4% paraformaldehyde, embedded in Histogel, dehydrated, and paraffin embedded using standard protocol. Five micron thick sections were cut and mounted on glass slides and immunostained for mouse SFTPC (1:1000), T1-α (15µg/ml), PDGFRα (1:50) α-smooth muscle actin (α-SMA; 1:500), Ki-67 (1:100), and Histone 2A.X (H2A.X; 1:100). Briefly, alveolosphere sections were deparaffinized in xylene and hydrated through ethanol series and water. Antigen retrieval was performed using citrate buffer at pH 6.0 in a 95°C water bath. Tissue sections were blocked using 5% normal goat or donkey serum and were then incubated in primary antibodies overnight at 4°C. Appropriate IgG isotype controls were also used to determine specificity of staining. Secondary detection was performed using anti-mouse Alexa Fluor 594/488-tagged secondary antibodies. Nuclei were counterstained with Hoechst 33342 dye for immunofluorescence detection. The stained alveolosphere sections were mounted in Vectashield and viewed and imaged in a Keyence BZ-X710 inverted microscope with brightfield as well as fluorescent imaging capability.

### Histochemical staining

β-galactosidase staining was performed on young and aged L-MSCs in culture, according to instructions provided in the Senescence Detection Kit (Abcam). Lipofuscin staining was performed on paraffin sections of alveolospheres following published protocol (Georgakopoulou et al., 2013).

### Cytokine Array

Cytokine array was carried out in the young and aged L-MSC culture media using Proteome Profiler Mouse XL Cytokine Array kit (R&D Systems) as per manufacturer ‘s instructions.

### Proteomics/Mass Spectrometry Analysis

Proteomics analysis was carried out following established protocols (Ludwig et al., 2016) with minor changes. Briefly, 5 ml of each cell culture media specimen were concentrated using Amicon Ultra 4ml, 3kDa molecular weight cut-off filters (Millipore) and protein concentrations were determined using Pierce BCA Protein Assay Kit. Proteins (10 µg) per sample were reduced with DTT and denatured at 70°C for 10 min prior to loading onto Novex NuPAGE 10% Bis-Tris Protein gels (Invitrogen, Cat. # NP0315BOX). The gels were stained overnight with Novex Colloidal Blue Staining kit (Invitrogen, Cat. # LC6025). Following de-staining, each lane were cut into three MW fractions and equilibrated in 100 mM ammonium bicarbonate (AmBc), each gel plug was then digested overnight with Trypsin Gold, Mass Spectrometry Grade (Promega, Cat. # V5280) following manufacturer ‘s instruction. Peptide digests (8µL each) were injected onto a 1260 Infinity nHPLC stack (Agilent Technologies), and separated using a 75 micron I.D. x 15 cm pulled tip C-18 column (Jupiter C-18 300 Å, 5 micron, Phenomenex). This system runs in-line with a Thermo Orbitrap Velos Pro hybrid mass spectrometer, equipped with a nano-electrospray source (Thermo Fisher Scientific), and all data were collected in CID mode. Searches were performed with a species specific subset of the UniprotKB database. The list of peptide IDs generated based on SEQUEST (Thermo Fisher Scientific) search results were filtered using Scaffold (Protein Sciences, Portland Oregon). Gene ontology assignments and pathway analysis were carried out using MetaCore (GeneGO Inc., St. Joseph, MI).

### Extracellular Flux Analysis

Analyses of cellular bioenergetics were performed using the Seahorse XFe96 Extracellular Flux Analyzer (Agilent, Santa Clara, CA). L-MSCs from young and aged mice were cultured for 24 hrs in low serum (1%) containing media. Cells were then detached and re-plated at a density of 25,000 cells /well into a XF96 microplate 5 hrs prior to assay in the same media. After the cells were attached, XF-DMEM media (DMEM supplemented with 5.5 mM glucose, 1 mM pyruvate and 4 mM L-Glutamine, pH 7.4 at 37 °C) was added to the wells, and cells were incubated for 1 hr prior to assay in a non-CO_2_ incubator at 37 °C. Mitochondrial stress test, a parallel measures of basal oxygen consumption rate (OCR) and extracellular acidification rate (ECAR), was carried out following sequential injections of oligomycin (1 µg/ml), carbonyl cyanide 4-(trifluoromethoxy) phenylhydrazone [FCCP (0.6 µM)], antimycin-A (10 µM), and 2-deoxyglucose (50 µM).

### ATP/ADP Ratio

This assay was performed using an ADP/ATP ratio assay kit (Abcam) following manufacturer ‘s instructions.

### H_2_O_2_-release Assay

This assay was performed based on protocol developed in our laboratory for quantitative measurement of extracellular H_2_O_2_release from adherent fibroblasts (Thannickal and Fanburg, 1995). This fluorimetric method relies on the oxidative conversion of homovanillic acid (HVA), a substituted phenolic compound, to its fluorescent dimer in the presence of H_2_O_2_and horseradish peroxidase (HRP). For this experiment, 150,000 young, aged, aged Nox4^-/+^ mouse lung L-MSCs were seeded in 6-well culture dishes. Next day, cells were serum starved (1%) overnight and treated with either vehicle or 2.5 ng/ml TGF-β1. Media was aspirated after 16 hrs post-treatment; cells were washed with 1 ml of HBSS (without calcium and magnesium) and incubated for 2 hrs in 1 ml of assay media containing 100 µM HVA and 5U/ml HRP in HBSS (containing calcium and magnesium). Reactions were stopped adding stop solution and fluorescence was measured in a BioTek microplate reader at excitation and emission maximum of 321 nm and 421 nm respectively. Extracellular H_2_O_2_ release was measured from standard curve generated from a known concentrations of H_2_O_2_. The data are normalized to cell number and H_2_O_2_ concentrations are presented as nanomoles/min/10^6^ L-MSCs.

### Quantitative RT-PCR

Total RNA was isolated from aged and aged Nox4^-/+^ L-MSCs using RNeasy® Mini Kit (Qiagen) and reverse transcribed using iScript Reverse Transcription SuperMix for RT-qPCR (Bio-Rad). Real-time PCR reactions were performed using SYBR® Green PCR Master Mix (Life Technologies) and gene specific primer pairs for Nox4 and β-actin (Integrated DNA Technologies; for primer sequences, see **Supplementary Table 6**). Reactions were carried out for 40 cycles (95°C for 15 sec, 60°C for 1 min) in a StepOnePlus Real Time PCR System (Life Technologies). Real-time PCR data (2^-ΔΔCt^) is presented as Nox4 mRNA expression normalized to β-actin.

### Targeted metabolomics

Concentrations of TCA metabolites in the young and aged L-MSCs were determined by Liquid chromatography tandem mass spectrometry (LCMS) following published protocols (Tan et al., 2014) with minor modifications. Dry stocks of analytically pure standards were weighed out and solubilized in MilliQ water and further diluted with 50% methanol to a 10 µg/ml stock. This stock was diluted to generate concentrations for standards ranging from 0.5 to 1000 ng/ml. Methanolic sample extracts were dried under nitrogen (N2) gas and reconstituted in 100 µl of 50% methanol. Methanolic samples and standards were derivatized by addition of 10 µl 0.1M O-benzylhydroxylamine hydrochloride (O-BHA) and 10 µl 0.25M N-(3-dimethylaminopropyl)-N′-ethylcarbodiimide hydrochloride (EDC) at RT for 1 hr. Samples were then transferred to a 13 x 100 mm borosilicate tube for liquid-liquid-extraction. Samples were then dried under N_2_ gas at RT and were reconstituted in 100 µl of 0.1% formic acid (FA) and transferred to HPLC vials for LCMS analysis. LCMS was carried out with a Prominence HPLC and API 4000 MS. 10 µl of sample was injected onto an Accucore 2.6 µm C18 100 x 2.1 mm column at 40°C for gradient separation. Column eluent was directed to the API 4000 MS operating system, controlled by Analyst 1.6.2 software. Post-acquisition data analysis was carried out using MultiQuant v3.0.3 software. All standard curve regressions were linear with 1/x^2^ weighting.

### Statistical Analysis

Unpaired Student ‘s T-test was used to compare between two experimental groups. When more than two experimental groups were present, data were analyzed by one-way Analysis of Variance (ANOVA). Tukey test was performed to compare between multiple groups. Data were tested for normality distribution by Shapiro-Wilk test before comparative analysis. Non-normal data (alveolosphere size; **Figure 1J** and **Figure 4I**) were natural log transformed and compared by one-way ANOVA and Tukey tests for multiple comparisons. Differences between the experimental groups were considered significant when *p < 0*.*05*.

## Acknowledgement

We thank Dr. Tingting Yuan and Dr. Stijn De Langhe for providing access to their laboratory for preparing paraffin blocks and cutting histological sections. We thank Dr. Robert Grabski and Mr. Shawn Williams for their assistance with confocal imaging at the High Resolution Imaging Facility at the University of Alabama at Birmingham (UAB). We also thank Dr. Stephen Barnes, Dr. Landon Wilson, and Mr. Taylor Berryhill at the Targeted Proteomics & Metabolomics Lab, UAB for their assistance with measuring TCA cycle metabolites. This work was supported by NIH grants, P01 HL114470 and R01 AG046210; and by VA Merit Award, I01BX003056 (to VJT). DC is supported by NIH/NHLBI grant, R01 HL152246 (to VJT); JAM is supported by NIH/NCI grant, P30 CA013148; JSD is supported by NIH grants, R01 HL128502 and P42 ES027723.

## Footnotes

### AUTHOR CONTRIBUTIONS

DC conceptualized the project, designed and conducted experiments, analyzed results and wrote the manuscript. RM assisted with experiments and graphical illustrations. SRS analyzed the cytokine array data and assisted with drawing the schematics. KGD assisted with alveolosphere assay. YW assisted with flow cytometry. KB assisted with hydrogen peroxide assay. DK assisted with generation of NOX4-/+ mouse line. VM counted and measured the alveolospheres using Image J. KK and JAM assisted with discovery proteomics on the cell culture samples and analyzed the mass spectrometry data. GB performed the Seahorse assay and analyzed the data. VDU assisted with Seahorse assay and edited the manuscript. JZ and JSD edited the manuscript. VJT conceived the project, designed experiments, analyzed data and co-wrote the manuscript.

### COMPETING INTERESTS STATEMENT

The authors declare no conflict of interest and there are no financial relationships with commercial entities which have interest in the subject of this manuscript.

## Notes

### Competing Interest Statement

The authors have declared no competing interest.

