## Supplementary Information for "Mesenchymal Stromal Cell Aging Impairs the Self-Organizing Capacity of Lung Alveolar Epithelial Stem Cells"

[1]Division of Pulmonary, Allergy, and Critical Care Medicine, Department of Medicine; [2]Department of Surgery; [3]Comprehensive Cancer Center Mass Spectrometry & Proteomics Shared Facility, [4]Department of Anesthesiology and Perioperative Medicine; [5]Department of Pathology; [6]Division of Preventive Medicine, Department of Medicine; University of Alabama at Birmingham, Birmingham, AL, USA; [7]John W. Deming Department of Medicine, Tulane University School of Medicine, New Orleans, Louisiana, USA

**Correspondence:**

Diptiman Chanda, Ph.D.
Assistant Professor of Medicine
Division of Pulmonary, Allergy, and Critical Care
Department of Medicine
University of Alabama at Birmingham
1900 University Blvd THT 541E, Birmingham, AL 35294-2180


**Or to:**

Victor J. Thannickal, M.D.
Professor and Harry B. Greenberg Chair
John W. Deming Department of Medicine
Tulane University School of Medicine
1430 Tulane Avenue, #8512
New Orleans, LA 70112


#### Figure supplement 1

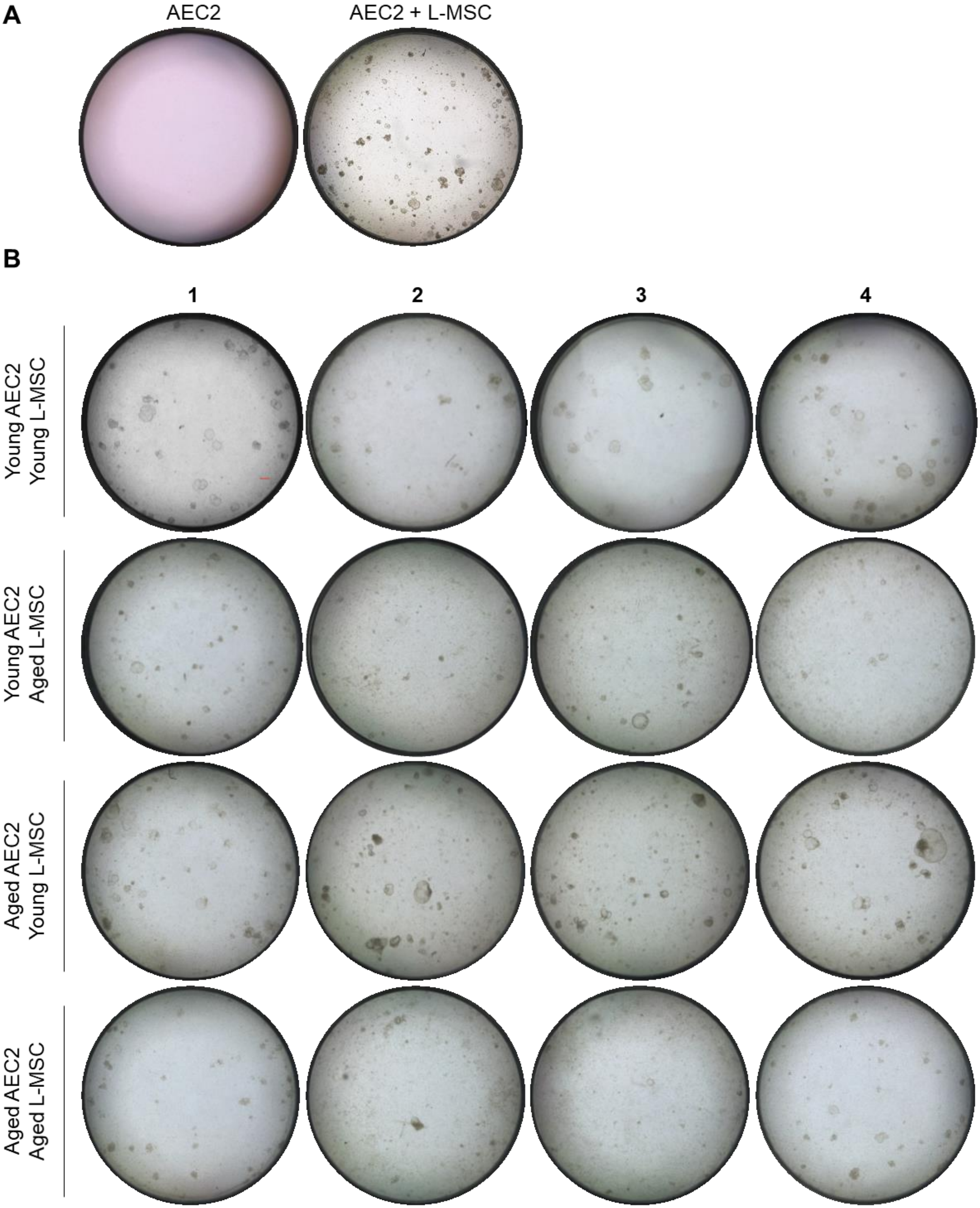

**Figure supplement 1:** (**A**) Type 2 alveolar epithelial cells (AEC2s) do not form alveolospheres in the absence of lung mesenchymal stromal cells (L-MSCs). (**B**) **Aging L-MSCs impair self-organization of AEC2s and alveolosphere formation** L-MSCs and AEC2s were isolated from young (3 months) and aged (24 months) mice and co-cultured as described. Varied combinations of AEC2s and L-MSCs from lungs of mice were studied (scale bar = 300 µm). Alveolospheres were imaged by brightfield microscopy after 12 days of co-culture. Experimental replicates are shown here.

### Figure supplement 2

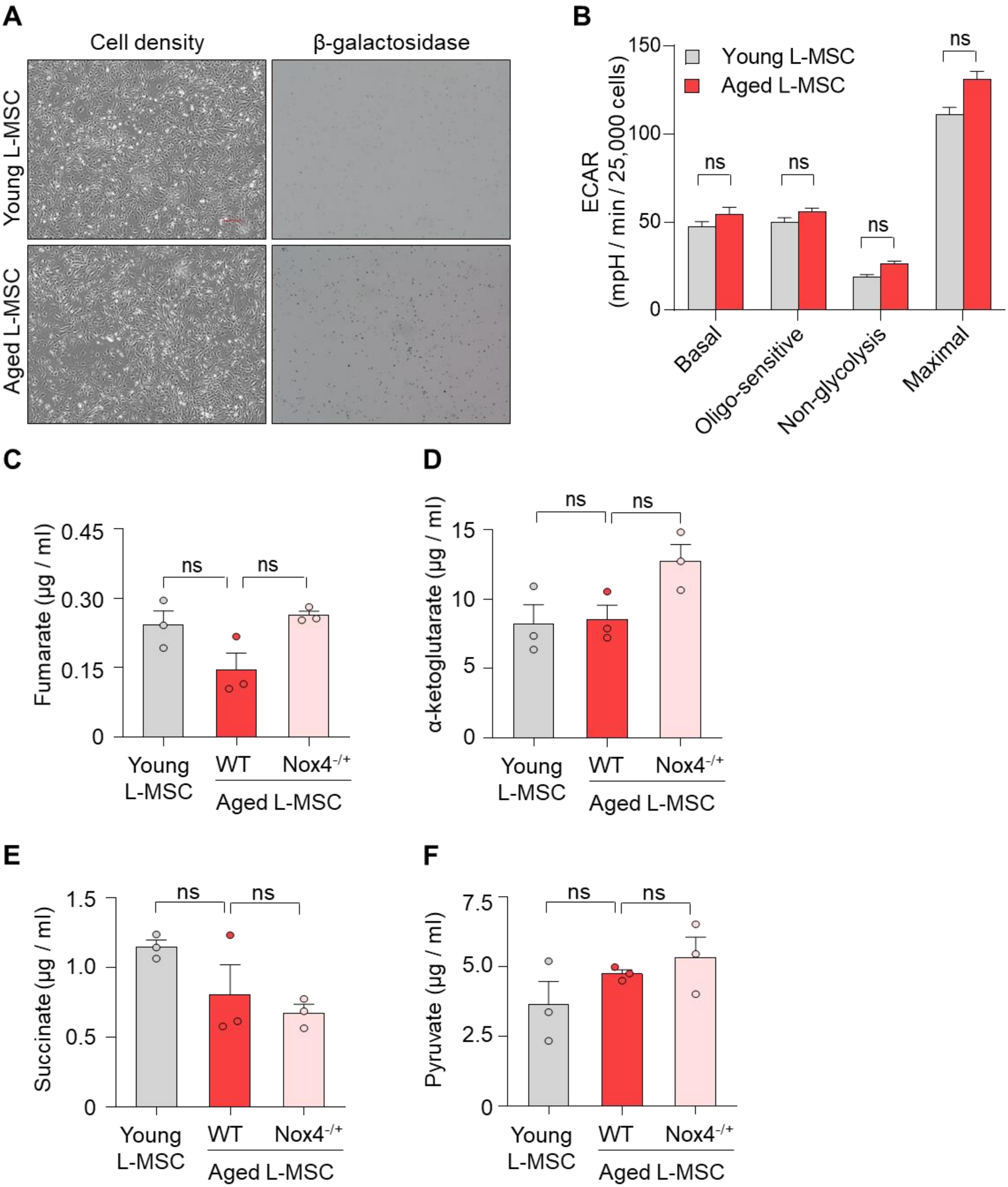

**<u>Figure supplement 2</u>**: **Aged L-MSCs show features of senescence and altered bioenergetics**. (**A**) β-galactosidase staining was performed to determine senescence of aged L-MSCs (vs. young) L-MSCs. Low magnification images are shown here (scale bar = 300 μm). (**B**) **Senescent L-MSCs demonstrate impaired bioenergetics**. Comparative extracellular acidification rates (ECAR) between young and aged L-MSCs are shown (n = 8; $p > 0.05$; ns = not significant). (**C-F**) Targeted metabolomics: Concentrations of fumarate, α-ketoglutarate, succinate, and pyruvate in young, aged, and Nox4-deficient L-MSCs were also determined (n = 3; $p > 0.05$; ns = not significant).

### Figure supplement 3

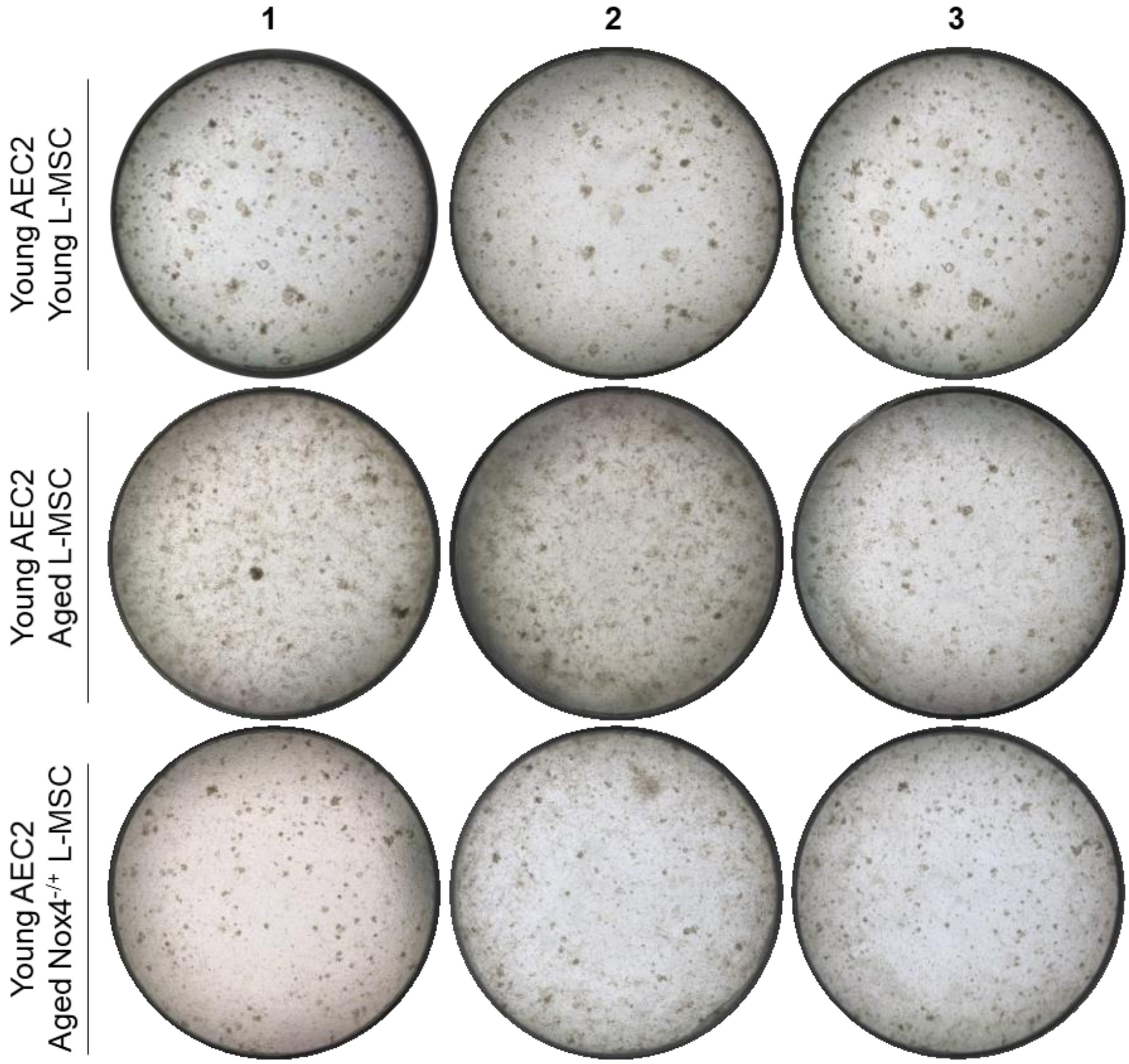


**Figure supplement 3: Nox4-deficiency in aged L-MSCs improves bioenergetics and restores AEC2 self-organization**. L-MScs were isolated from young, aged, and $Nox4^{-/+}$ aged mice and co-cultured with young AEC2s as described earlier. Varied combinations of AEC2s and L-MSCs from lungs of mice were studied (scale bar = 300 µm). Alveolospheres were imaged by brightfield microscopy after 12 days of co-culture. Experimental replicates are shown here.

#### Table supplement 1: Top Proteins Changes (Aged *vs.* Young mouse L-MSCs)

| UniProtKB Name | Gene Name | UProt Acc# | GeneID | NetworkID | GO Function/ Biological Processes | Old(avg) | Y(avg) | SAM | Ttest | Fold(O/Y) |
|---|---|---|---|---|---|---|---|---|---|---|
| Procollagen-lysine,2-oxoglutarate 5-dioxygenase 1 | procollagen-lysine, 2-oxoglutarate 5-dioxygenase 1(Plod1) | Q9R0E2 | 18822 | PLOD1 | *** oxidoreductase/ $H_2O_2$ catabolic/ response to hypoxia/ metal binding/ peptidyl-lysine hydroxylation** | 5.38 | 1.61 | 1.32 | 0.0340 | 3.34 |
| Peroxidasin | peroxidasin(Pxdn) | Q3UQ28 | 69675 | PXDN | *** oxidoreductase/ $H_2O_2$ catabolic/ metal-heme binding/ peroxidase/ response to ox-stress** | 9.56 | 3.63 | 1.08 | 0.0265 | 2.63 |
| Integrin beta-2 | integrin beta 2(Itgb2) | P11835 | 16414 | Integrin | *** HSP-binding/ pos reg NO process/ cell-cell adhesion** | 2.62 | 0.00 | - | - | 2.62 |
| Keratin, type I cytoskeletal 17 | keratin 17(Krt17) | Q9QWL7 | 16667 | Keratin 17 | intermediate filamate organisation/ pos cell growth | 6.73 | 2.79 | 0.81 | 0.0452 | 2.41 |
| Legumain | legumain(Lgmn) | O89017 | 19141 | Legumain | protease/ endopeptidase | 5.62 | 2.54 | 0.79 | 0.1044 | 2.21 |
| Keratin, type II cytoskeletal 2 epidermal | keratin 2(Krt2) | Q3TTY5 | 16681 | Keratin 2 | cytoskeletal protein binding/ keratinocyte development | 7.90 | 3.69 | 0.93 | 0.0228 | 2.14 |
| Beta-galactosidase | galactosidase, beta 1(Glb1) | P23780 | 12091 | BGAL | *** hydrolase activity/ galactose catabolism/ proton donor** | 2.09 | 0.00 | - | - | 2.09 |
| Plastin-3 | plastin 3 (T-isoform)(Pls3) | Q99K51 | 102866 | T-plastin | actin filiment binding/ | 9.90 | 4.81 | 0.79 | 0.0543 | 2.06 |
| ATP-binding cassette sub-family A member 8-B | ATP-binding cassette, sub-family A (ABC1) | Q8K440 | 27404 | ABCA8 | ATPase activity/ lipid trasport | 4.08 | 2.03 | 0.83 | 0.0595 | 2.01 |
| Collagen alpha-1(XII) chain | collagen, type XII, alpha 1(Col12a1) | Q60847 | 12816 | Collagen XII | hydroxylation/ cell adhesion | 52.29 | 29.03 | 1.16 | 0.0144 | 1.80 |
| Aspartyl aminopeptidase | aspartyl aminopeptidase(Dnpep) | Q9Z2W0 | 13437 | Dnpep | metaloprotease/ Zn binding | 3.79 | 2.12 | 1.22 | 0.0125 | 1.79 |
| ATP synthase subunit alpha, mitochondrial | ATP synthase, H+ transporting, mitochondrial F1 complex, alpha subunit 1(Atp5a1) | Q03265 | 11946 | ATP5A | ATP metabolic process/ protease binding | 1.63 | 0.00 | - | - | 1.63 |
| Alpha-enolase | enolase 1, alpha non-neuron(Eno1) | P17182 | 13806 | ENO | *** proton donor/ glycolysis/ Mg binding** | 24.23 | 15.88 | 0.96 | 0.0561 | 1.53 |
| Threonine--tRNA ligase, cytoplasmic | threonyl-tRNA synthetase(Tars) | Q9D0R2 | 110960 | SYTC | Zn binding/ threonyl-tRNA aminoacylation | 0.00 | 1.46 | - | - | -1.46 |
| Matrix Gla protein | matrix Gla protein(Mgp) | P19788 | 17313 | MGP | lung development/ Ca binding | 6.81 | 11.23 | 0.95 | 0.0359 | -1.65 |
| NADP-dependent malic enzyme | malic enzyme 1, NADP(+)-dependent, cytosolic(Me1) | P06801 | 17436 | ME1 | *** oxidoreductase/ proton donor** | 1.88 | 3.13 | 0.90 | 0.0555 | -1.67 |
| Elongation factor 1-alpha 1 | eukaryotic translation elongation factor 1 alpha 1(Eef1a1) | P10126 | 13627 | eEF1A | *** GTPase activity/ kinase binding/ neg reg reactive ox species** | 5.14 | 8.88 | 0.90 | 0.0419 | -1.73 |
| Lysosome-associated membrane glycoprotein 2 | lysosomal-associated membrane protein 2(Lamp2) | P17047 | 16784 | LAMP2 | pos autophagy/ reg protein stability | 0.00 | 1.74 | - | - | -1.74 |
| Peroxiredoxin-1 | peroxiredoxin 1(Prdx1) | P35700 | 18477 | Peroxiredoxin | *** oxidoreductase/ peroxidase/ heme binding/ redox homeostatis** | 8.09 | 14.25 | 1.05 | 0.0133 | -1.76 |
| Complement component 1, r subcomponent | complement component 1, r subcomponent A(C1ra) | Q566I6 | 50909 | C1RA | serine protease/ hydroxylation/ Ca binding/ complement activation | 2.85 | 5.46 | 1.31 | 0.0141 | -1.92 |
| Latent-transforming growth factor beta-binding protein 1 | latent transforming growth factor beta binding protein 1(Ltbp1) | Q8CG19 | 268977 | LTBP1 | Ca binding/ hydroxylation/ TGFbeta binding | 2.19 | 4.20 | 1.42 | 0.0272 | -1.92 |
| Stromal cell-derived factor 1 | chemokine (C-X-C motif) ligand 12(Cxcl12) | P40224 | 20315 | SDF-1 | CXC chemokine activity/ defense response/ cell-cell adhesion | 1.45 | 2.81 | 1.19 | 0.0267 | -1.94 |
| Dickkopf-related protein 3 | dickkopf WNT signaling pathway inhibitor 3(Dkk3) | Q9QUN9 | 50781 | DKK3 | neg reg Wnt signaling | 1.86 | 3.66 | 1.18 | 0.0133 | -1.97 |
| Peptidyl-prolyl cis-trans isomerase FKBP10 | FK506 binding protein 10(Fkbp10) | Q61576 | 14230 | FKBP10 | Ca binding/ peptidyl-proline modification | 2.16 | 4.27 | 1.59 | 0.0202 | -1.98 |
| Elastin | elastin(Eln) | P54320 | 13717 | Elastin | hydroxylation/ ECM binding & organization | 0.00 | 2.08 | - | - | -2.08 |
| L-lactate dehydrogenase A chain | lactate dehydrogenase A(Ldha) | P06151 | 16828 | LDHA | *** oxidoreductase/ LDH activity** | 6.61 | 13.87 | 0.78 | 0.0431 | -2.10 |
| Nucleoside diphosphate kinase B | NME/NM23 nucleoside diphosphate kinase 2(Nme2) | Q01768 | 18103 | NDPK B | *** kinase acitivity/ ATP binding/ response to ox-stress** | 5.61 | 12.20 | 0.97 | 0.0289 | -2.17 |
| C-type mannose receptor 2 | mannose receptor, C type 2(Mrc2) | Q64449 | 17534 | ENDO180 | collagen binding & catabolism process/ endocytosis | 1.43 | 3.34 | 0.83 | 0.0841 | -2.33 |
| 14-3-3 protein epsilon | tyrosine 3-monooxygenase/tryptophan 5-monooxygenase activation protein, epsilon | D6REF3 | 22627 | 14-3-3 | *** response to stress/ oxidoreductase/ cell cycle checkpoint associated** | 1.44 | 3.37 | 2.46 | 0.0185 | -2.34 |
| Insulin-like growth factor-binding protein 4 | insulin-like growth factor binding protein 4(Igfbp4) | P47879 | 16010 | IBP | regulation of cell growth/ glucose metabolism/ infammation response | 2.58 | 6.43 | 0.81 | 0.0761 | -2.50 |
| Ecm1 protein | extracellular matrix protein 1(Ecm1) | B7ZNR0 | 13601 | ECM1 | signal transduction | 2.95 | 7.41 | 1.12 | 0.0779 | -2.51 |
| Glyceraldehyde-3-phosphate dehydrogenase | glyceraldehyde-3-phosphate dehydrogenase(Gapdh) | P16858 | 14433 | G3P2 | *** oxidoreductase/ glycolysis/ aspartate endopeptidase inhibition/** | 3.43 | 8.68 | 1.27 | 0.0057 | -2.53 |
| metalloproteinase with thrombospondin motifs 2 | metallopeptidase (reprolysin type) with thrombospondin type 1 motif, 2(Adamts2) | Q8C9W3 | 216725 | ADAM-TS2 | metaloprotease/ Zn binding/ lung development | 1.35 | 3.62 | 0.87 | 0.0958 | -2.68 |
| Mimecan | osteoglycin(Ogn) | Q62000 | 18295 | Osteoglycin | growth factor/ TGF-beta associated | 2.98 | 8.64 | 1.39 | 0.0347 | -2.90 |
| Stromelysin-1 | matrix metallopeptidase 3(Mmp3) | P28862 | 17392 | Stromelysin-1 | metaloprotease/ Ca & Zn binding/ hydrolase | 6.64 | 20.92 | 1.46 | 0.0259 | -3.15 |
| Gelsolin | gelsolin(Gsn) | P13020 | 227753 | Gelsolin | Ca binding/ actin associated/ aging processes | 19.90 | 67.30 | 2.80 | 0.0039 | -3.38 |

Note: *GO associations were derived from UniProtKB and DAVID GO databases*

**** oxidative-stress/ redox associated***

Table supplement 2: Enrichment of proteins by Gene Ontology Localizations

**Enrichment by GO Localizations**

| # | Localizations | Total | p-value | FDR | In Data | Network Objects from Active Data |
|---|---|---|---|---|---|---|
| 1 | cytoplasmic, known to go to extracellular space | 4015 | 7.9E-18 | 2.2E-15 | 31 | PLOD1, SDF-1, eEF1A1, MGP, BGAL, ADAM-TS2, G3P2, Keratin 17, LAMP2, IBP4, Elastin, DKK3, ITGB2, Gelsolin, ATP5A, LDHA, PRDX1, ECM1, 14-3-3 epsilon, LTBP1, SYTC, Osteoglycin, ENO1, Collagen XII, Dnpep, PXDN, NDPK B, Legumain, Stromelysin-1, Keratin 2, C1RA |
| 2 | extracellular exosome | 2237 | 5.9E-15 | 4.5E-13 | 23 | PLOD1, SDF-1, eEF1A1, MGP, BGAL, G3P2, Keratin 17, LAMP2, ITGB2, Gelsolin, ATP5A, LDHA, PRDX1, ECM1, 14-3-3 epsilon, SYTC, Osteoglycin, ENO1, Collagen XII, PXDN, NDPK B, Legumain, Keratin 2 |
| 3 | collagen-containing extracellular matrix | 529 | 4.6E-09 | 1.8E-07 | 10 | PLOD1, SDF-1, MGP, ADAM-TS2, Elastin, ECM1, LTBP1, Osteoglycin, Collagen XII, PXDN |
| 4 | extracellular matrix | 740 | 8.4E-09 | 2.6E-07 | 11 | PLOD1, SDF-1, MGP, ADAM-TS2, Elastin, ECM1, LTBP1, Osteoglycin, Collagen XII, PXDN, Stromelysin-1 |
| 5 | myelin sheath | 221 | 3.7E-08 | 1.0E-06 | 7 | eEF1A1, G3P2, Gelsolin, ATP5A, PRDX1, ENO1, NDPK B |
| 6 | lysosomal lumen | 101 | 1.6E-05 | 4.0E-04 | 4 | BGAL, LAMP2, Osteoglycin, Legumain |
| 7 | ficolin-1-rich granule lumen | 129 | 4.2E-05 | 8.2E-04 | 4 | eEF1A1, BGAL, Gelsolin, NDPK B |
| 8 | extracellular matrix component | 65 | 1.3E-04 | 2.2E-03 | 3 | Elastin, LTBP1, Collagen XII |
| 9 | secretory granule lumen | 336 | 1.4E-04 | 2.2E-03 | 5 | eEF1A1, BGAL, Gelsolin, ECM1, NDPK B |
| 10 | vacuolar lumen | 180 | 1.5E-04 | 2.2E-03 | 4 | BGAL, LAMP2, Osteoglycin, Legumain |
| 11 | vesicle lumen | 342 | 1.5E-04 | 2.2E-03 | 5 | eEF1A1, BGAL, Gelsolin, ECM1, NDPK B |
| 12 | mitochondrion | 2183 | 2.8E-04 | 3.9E-03 | 11 | G3P2, Elastin, ABCA8, ATP5A, LDHA, PRDX1, 14-3-3 epsilon, LTBP1, NDPK B, Stromelysin-1, ME1 |
| 13 | perinuclear region of cytoplasm | 884 | 3.1E-04 | 4.1E-03 | 7 | BGAL, G3P2, LAMP2, Gelsolin, LTBP1, NDPK B, Legumain |
| 14 | cell-substrate adherens junction | 433 | 4.6E-04 | 5.5E-03 | 5 | ITGB2, Gelsolin, 14-3-3 epsilon, ENDO180, NDPK B |
| 15 | secretory granule | 1002 | 6.6E-04 | 7.0E-03 | 7 | eEF1A1, BGAL, LAMP2, ITGB2, Gelsolin, ECM1, NDPK B |
| 16 | cytoplasm | 19040 | 8.9E-04 | 9.1E-03 | 38 | PLOD1, SDF-1, eEF1A1, MGP, eEF1A, Peroxiredoxin, BGAL, G3P2, Keratin 17, T-plastin, LAMP2, IBP4, Elastin, ABCA8, ITGB2, Gelsolin, ATP5A, IBP, LDHA, PRDX1, ECM1, 14-3-3 epsilon, FKBP10, ENO, LTBP1, SYTC, Osteoglycin, ENO1, Collagen XII, Dnpep, LAMP2-C, PXDN, 14-3-3, NDPK B, Legumain, Stromelysin-1, ME1, Keratin 2 |
| 17 | large latent transforming growth factor-beta complex | 1 | 1.5E-03 | 1.4E-02 | 1 | LTBP1 |
| 18 | collagen type XII trimer | 1 | 1.5E-03 | 1.4E-02 | 1 | Collagen XII |
| 19 | adherens junction | 581 | 1.7E-03 | 1.4E-02 | 5 | ITGB2, Gelsolin, 14-3-3 epsilon, ENDO180, NDPK B |
| 20 | supramolecular fiber | 1179 | 1.7E-03 | 1.4E-02 | 7 | Keratin 17, T-plastin, Elastin, LTBP1, ENO1, NDPK B, Keratin 2 |
| 21 | endomembrane system | 5557 | 1.7E-03 | 1.4E-02 | 17 | PLOD1, eEF1A1, MGP, BGAL, G3P2, LAMP2, IBP4, ITGB2, Gelsolin, ECM1, FKBP10, LTBP1, Osteoglycin, Collagen XII, PXDN, NDPK B, Legumain |
| 22 | supramolecular complex | 1187 | 1.8E-03 | 1.4E-02 | 7 | Keratin 17, T-plastin, Elastin, LTBP1, ENO1, NDPK B, Keratin 2 |
| 23 | endoplasmic reticulum lumen | 347 | 1.8E-03 | 1.4E-02 | 4 | IBP4, FKBP10, LTBP1, Collagen XII |
| 24 | anchoring junction | 604 | 2.0E-03 | 1.5E-02 | 5 | ITGB2, Gelsolin, 14-3-3 epsilon, ENDO180, NDPK B |
| 25 | secretory vesicle | 1232 | 2.2E-03 | 1.6E-02 | 7 | eEF1A1, BGAL, LAMP2, ITGB2, Gelsolin, ECM1, NDPK B |
| 26 | vacuole | 939 | 2.6E-03 | 1.8E-02 | 6 | eEF1A1, BGAL, LAMP2, Osteoglycin, Dnpep, Legumain |
| 27 | integrin alphaL/M-beta2 complex | 2 | 3.0E-03 | 1.8E-02 | 1 | ITGB2 |
| 28 | ruffle | 210 | 3.8E-03 | 2.3E-02 | 3 | eEF1A1, Gelsolin, NDPK B |
| 29 | membrane raft | 433 | 4.0E-03 | 2.3E-02 | 4 | LAMP2, ITGB2, ATP5A, ENO1 |
| 30 | cytoplasmic vesicle part | 1776 | 4.4E-03 | 2.4E-02 | 8 | eEF1A1, BGAL, LAMP2, ITGB2, Gelsolin, ECM1, NDPK B, Legumain |

#### Table supplement 3: Enrichment of proteins by Gene Ontology Processes

Enrichment by GO Processes

| # | Processes | Total | p-value | FDR | In Data | Network Objects from Active Data |
|---|---|---|---|---|---|---|
| 1 | negative regulation of reactive oxygen species metabolic process | 102 | 1.3E-06 | 1.0E-03 | 4 | eEF1A1, eEF1A, Peroxiredoxin, Stromelysin-1 |
| 2 | extracellular matrix organization | 466 | 1.5E-06 | 1.0E-03 | 6 | PLOD1, Peroxiredoxin, Collagen XII, PXDN, Gelsolin, Stromelysin-1 |
| 3 | extracellular structure organization | 540 | 3.5E-06 | 1.6E-03 | 6 | PLOD1, Peroxiredoxin, Collagen XII, PXDN, Gelsolin, Stromelysin-1 |
| 4 | cellular component organization | 7470 | 7.8E-06 | 2.2E-03 | 16 | PLOD1, SDF-1, eEF1A1, FKBP10, MGP, eEF1A, Osteoglycin, Peroxiredoxin, Collagen XII, G3P2, Keratin 17, PXDN, NDPK B, Gelsolin, Stromelysin-1, Keratin 2 |
| 5 | hydrogen peroxide catabolic process | 48 | 8.1E-06 | 2.2E-03 | 3 | Peroxiredoxin, PXDN, PRDX1 |
| 6 | positive regulation of epidermis development | 51 | 9.7E-06 | 2.2E-03 | 3 | Keratin 17, NDPK B, Keratin 2 |
| 7 | cellular component organization or biogenesis | 7737 | 1.3E-05 | 2.4E-03 | 16 | PLOD1, SDF-1, eEF1A1, FKBP10, MGP, eEF1A, Osteoglycin, Peroxiredoxin, Collagen XII, G3P2, Keratin 17, PXDN, NDPK B, Gelsolin, Stromelysin-1, Keratin 2 |
| 8 | anatomical structure development | 7828 | 1.5E-05 | 2.4E-03 | 16 | PLOD1, SDF-1, eEF1A1, MGP, LTBP1, eEF1A, Osteoglycin, Peroxiredoxin, Collagen XII, G3P2, Keratin 17, NDPK B, DKK3, Gelsolin, Keratin 2, PRDX1 |
| 9 | cell killing | 193 | 1.7E-05 | 2.4E-03 | 4 | SDF-1, Peroxiredoxin, G3P2, PRDX1 |
| 10 | cell activation | 1558 | 1.8E-05 | 2.4E-03 | 8 | SDF-1, eEF1A1, eEF1A, Peroxiredoxin, NDPK B, Gelsolin, Keratin 2, PRDX1 |
| 11 | regulation of chaperone-mediated autophagy | 9 | 2.3E-05 | 2.7E-03 | 2 | eEF1A1, eEF1A |
| 12 | hydrogen peroxide metabolic process | 69 | 2.4E-05 | 2.7E-03 | 3 | Peroxiredoxin, PXDN, PRDX1 |
| 13 | developmental process | 8282 | 3.4E-05 | 3.6E-03 | 16 | PLOD1, SDF-1, eEF1A1, MGP, LTBP1, eEF1A, Osteoglycin, Peroxiredoxin, Collagen XII, G3P2, Keratin 17, NDPK B, DKK3, Gelsolin, Keratin 2, PRDX1 |
| 14 | cellular response to chemical stimulus | 4503 | 4.6E-05 | 4.0E-03 | 12 | PLOD1, SDF-1, eEF1A1, LTBP1, eEF1A, Peroxiredoxin, G3P2, PXDN, NDPK B, Gelsolin, Stromelysin-1, PRDX1 |
| 15 | antibiotic catabolic process | 86 | 4.7E-05 | 4.0E-03 | 3 | Peroxiredoxin, PXDN, PRDX1 |
| 16 | multicellular organism development | 7341 | 4.7E-05 | 4.0E-03 | 15 | SDF-1, eEF1A1, MGP, LTBP1, eEF1A, Osteoglycin, Peroxiredoxin, Collagen XII, G3P2, Keratin 17, NDPK B, DKK3, Gelsolin, Keratin 2, PRDX1 |
| 17 | leukocyte activation | 1334 | 6.1E-05 | 4.9E-03 | 7 | SDF-1, eEF1A1, eEF1A, Peroxiredoxin, NDPK B, Gelsolin, PRDX1 |
| 18 | system development | 6534 | 7.1E-05 | 5.2E-03 | 14 | SDF-1, eEF1A1, MGP, LTBP1, eEF1A, Osteoglycin, Peroxiredoxin, Collagen XII, Keratin 17, NDPK B, DKK3, Gelsolin, Keratin 2, PRDX1 |
| 19 | collagen metabolic process | 100 | 7.3E-05 | 5.2E-03 | 3 | PLOD1, Collagen XII, Stromelysin-1 |
| 20 | cofactor catabolic process | 104 | 8.2E-05 | 5.4E-03 | 3 | Peroxiredoxin, PXDN, PRDX1 |
| 21 | regulation of reactive oxygen species metabolic process | 291 | 8.4E-05 | 5.4E-03 | 4 | eEF1A1, eEF1A, Peroxiredoxin, Stromelysin-1 |
| 22 | removal of superoxide radicals | 19 | 1.1E-04 | 6.4E-03 | 2 | Peroxiredoxin, PRDX1 |
| 23 | regulation of epidermis development | 115 | 1.1E-04 | 6.4E-03 | 3 | Keratin 17, NDPK B, Keratin 2 |
| 24 | neutrophil degranulation | 624 | 1.2E-04 | 6.4E-03 | 5 | eEF1A1, eEF1A, Peroxiredoxin, NDPK B, Gelsolin |
| 25 | neutrophil activation involved in immune response | 628 | 1.3E-04 | 6.4E-03 | 5 | eEF1A1, eEF1A, Peroxiredoxin, NDPK B, Gelsolin |
| 26 | immune response | 2684 | 1.3E-04 | 6.4E-03 | 9 | SDF-1, eEF1A1, eEF1A, Peroxiredoxin, G3P2, PXDN, NDPK B, Gelsolin, PRDX1 |
| 27 | supramolecular fiber organization | 632 | 1.3E-04 | 6.4E-03 | 5 | PLOD1, Collagen XII, Keratin 17, Gelsolin, Keratin 2 |
| 28 | neutrophil activation | 637 | 1.4E-04 | 6.4E-03 | 5 | eEF1A1, eEF1A, Peroxiredoxin, NDPK B, Gelsolin |
| 29 | granulocyte activation | 642 | 1.4E-04 | 6.4E-03 | 5 | eEF1A1, eEF1A, Peroxiredoxin, NDPK B, Gelsolin |
| 30 | neutrophil mediated immunity | 644 | 1.4E-04 | 6.4E-03 | 5 | eEF1A1, eEF1A, Peroxiredoxin, NDPK B, Gelsolin |
| 31 | cellular oxidant detoxification | 126 | 1.5E-04 | 6.4E-03 | 3 | Peroxiredoxin, PXDN, PRDX1 |
| 32 | leukocyte mediated immunity | 1054 | 1.5E-04 | 6.4E-03 | 6 | eEF1A1, eEF1A, Peroxiredoxin, NDPK B, Gelsolin, PRDX1 |
| 33 | leukocyte degranulation | 658 | 1.6E-04 | 6.5E-03 | 5 | eEF1A1, eEF1A, Peroxiredoxin, NDPK B, Gelsolin |
| 34 | cellular detoxification | 131 | 1.6E-04 | 6.5E-03 | 3 | Peroxiredoxin, PXDN, PRDX1 |
| 35 | cellular response to superoxide | 24 | 1.8E-04 | 6.6E-03 | 2 | Peroxiredoxin, PRDX1 |
| 36 | cellular response to oxygen radical | 24 | 1.8E-04 | 6.6E-03 | 2 | Peroxiredoxin, PRDX1 |
| 37 | response to oxidative stress | 676 | 1.8E-04 | 6.6E-03 | 5 | Peroxiredoxin, PXDN, NDPK B, Stromelysin-1, PRDX1 |
| 38 | myeloid cell activation involved in immune response | 683 | 1.9E-04 | 6.7E-03 | 5 | eEF1A1, eEF1A, Peroxiredoxin, NDPK B, Gelsolin |
| 39 | myeloid leukocyte mediated immunity | 689 | 2.0E-04 | 6.7E-03 | 5 | eEF1A1, eEF1A, Peroxiredoxin, NDPK B, Gelsolin |
| 40 | response to inorganic substance | 1108 | 2.0E-04 | 6.7E-03 | 6 | MGP, eEF1A, Peroxiredoxin, Gelsolin, Stromelysin-1, PRDX1 |
| 41 | response to superoxide | 28 | 2.4E-04 | 8.0E-03 | 2 | Peroxiredoxin, PRDX1 |
| 42 | cellular response to oxidative stress | 394 | 2.7E-04 | 8.7E-03 | 4 | Peroxiredoxin, NDPK B, Stromelysin-1, PRDX1 |
| 43 | detoxification | 159 | 2.9E-04 | 9.1E-03 | 3 | Peroxiredoxin, PXDN, PRDX1 |
| 44 | response to oxygen radical | 31 | 3.0E-04 | 9.1E-03 | 2 | Peroxiredoxin, PRDX1 |
| 45 | intermediate filament organization | 32 | 3.1E-04 | 9.3E-03 | 2 | Keratin 17, Keratin 2 |
| 46 | regulation of hydrogen peroxide metabolic process | 32 | 3.1E-04 | 9.3E-03 | 2 | Peroxiredoxin, Stromelysin-1 |
| 47 | reactive oxygen species metabolic process | 167 | 3.3E-04 | 9.7E-03 | 3 | Peroxiredoxin, PXDN, PRDX1 |
| 48 | myeloid leukocyte activation | 780 | 3.5E-04 | 9.9E-03 | 5 | eEF1A1, eEF1A, Peroxiredoxin, NDPK B, Gelsolin |
| 49 | leukocyte activation involved in immune response | 833 | 4.7E-04 | 1.3E-02 | 5 | eEF1A1, eEF1A, Peroxiredoxin, NDPK B, Gelsolin |
| 50 | cell activation involved in immune response | 837 | 4.8E-04 | 1.3E-02 | 5 | eEF1A1, eEF1A, Peroxiredoxin, NDPK B, Gelsolin |

Table supplement 4: Top Toxic Pathologies

Enrichment by Toxic Pathologies

| # | Toxic pathologies | Total | p-value | FDR | In Data | Network Objects from Active Data |
|---|---|---|---|---|---|---|
| 1 | Lung-fibrosis | 396 | 1.4E-03 | 4.6E-01 | 8 | SDF-1, Peroxiredoxin, Elastin, LDHA, PRDX1, FKBP10, Integrin, Stromelysin-1 |
| 2 | Hippocampus-apoptosis | 281 | 5.0E-03 | 4.8E-01 | 6 | Peroxiredoxin, 14-3-3 epsilon, ENO, ENO1, Integrin, 14-3-3 |
| 3 | Brain lesions | 530 | 9.0E-03 | 6.4E-01 | 8 | SDF-1, Peroxiredoxin, ATP5A, 14-3-3 epsilon, ENO, ENO1, Integrin, 14-3-3 |
| 4 | Intestine-proliferation | 367 | 1.8E-02 | 6.4E-01 | 6 | IBP, LDHA, Integrin, NDPK B, Stromelysin-1, ME1 |
| 5 | Brain-lipid peroxidation | 489 | 2.0E-02 | 6.4E-01 | 7 | Peroxiredoxin, ATP5A, 14-3-3 epsilon, ENO, ENO1, Integrin, 14-3-3 |
| 6 | Lung-inflammation | 390 | 2.3E-02 | 6.4E-01 | 6 | SDF-1, Peroxiredoxin, Elastin, PRDX1, Integrin, Stromelysin-1 |
| 7 | Lung pathology | 920 | 3.0E-02 | 6.4E-01 | 10 | SDF-1, Peroxiredoxin, Elastin, IBP, LDHA, PRDX1, FKBP10, ENO, Integrin, Stromelysin-1 |
| 8 | Brain-degeneration | 426 | 3.5E-02 | 6.4E-01 | 6 | Peroxiredoxin, 14-3-3 epsilon, ENO, ENO1, Integrin, 14-3-3 |

Table supplement 5: Top 10 Networks

Network List

| # | Network | GO processes | Total nodes | Seed nodes | p-Value | zScore | gScore |
|---|---|---|---|---|---|---|---|
| 1 | c-Myc, Oct-3/4, SOX9, MITF, Cyclin D1 | canonical Wnt signaling pathway (23.5%; 1.510e-21), cell-cell signaling by wnt (29.4%; 1.362e-17), Wnt signaling pathway (29.4%; 1.362e-17), regulation of canonical Wnt signaling pathway (26.5%; 3.388e-17), cell surface receptor signaling pathway involved in cell-cell signaling (30.9%; 5.320e-17) | 68 | 36 | 3.8E-128 | 3.5E+02 | 350.13 |
| 2 | NDPK B, p53, ETS1, Rb protein, IGF-1 | regulation of cell population proliferation (65.2%; 6.772e-36), negative regulation of cell population proliferation (47.8%; 1.135e-33), regulation of cell death (52.2%; 1.828e-22), negative regulation of cellular process (76.1%; 4.205e-22), response to endogenous stimulus (52.2%; 9.625e-22) | 95 | 36 | 3.0E-121 | 3.0E+02 | 296.21 |
| 3 | NDPK B, TGF-beta 1, MNT, GAS1, MAD4 | regulation of cell population proliferation (58.8%; 1.719e-25), cellular response to organic substance (67.5%; 1.209e-24), response to organic substance (73.8%; 4.263e-24), cellular response to chemical stimulus (71.2%; 8.126e-24), negative regulation of cell population proliferation (41.2%; 5.144e-23) | 80 | 34 | 6.9E-116 | 3.0E+02 | 304.86 |
| 4 | Elk-1, FOXO3A, CDC25C, CDK7, Bax | cell cycle process (51.8%; 3.296e-41), cell cycle (54.5%; 1.619e-38), mitotic cell cycle process (39.1%; 4.211e-34), mitotic cell cycle (41.8%; 1.276e-33), regulation of cell cycle process (40.9%; 8.285e-32) | 111 | 33 | 7.8E-106 | 2.5E+02 | 251.17 |
| 5 | PRDX1, Oxidized thioredoxin, Thioredoxin, Sirtuin1, KEAP1 | response to endogenous stimulus (68.5%; 2.135e-30), response to hormone (58.9%; 3.692e-29), response to oxygen-containing compound (65.8%; 2.108e-26), response to organic substance (78.1%; 1.624e-25), cellular response to oxygen-containing compound (56.2%; 1.625e-25) | 88 | 28 | 1.3E-90 | 2.5E+02 | 247.96 |
| 6 | G3P2, SIAH1, Norepinephrine extracellular region, IGF-1, Insulin processed | response to hormone (62.2%; 2.238e-20), response to peptide (48.9%; 3.392e-19), cellular response to oxygen-containing compound (60.0%; 2.686e-18), response to endogenous stimulus (66.7%; 2.956e-18), response to oxygen-containing compound (66.7%; 2.906e-17) | 53 | 21 | 1.9E-69 | 2.4E+02 | 238.16 |

**Table supplement 6: Key Resources**

| Reagent or Resources | Source | Identifier |
|---|---|---|
| **Mouse reactive antibodies used for flow cytometry** | | |
| Podoplanin (T1-α) monoclonal PE-Cyanine7 | eBioscience/ ThermoFisher Scientific | 25-5381-80 |
| CD31 (PECAM-1) FITC | eBioscience/ ThermoFisher Scientific | 11-0311-82 |
| CD326 (EpCAM) monoclonal APC-eFluor780 | eBioscience/ ThermoFisher Scientific | 47-5791-82 |
| CD24 Monoclonal, APC | eBioscience/ ThermoFisher Scientific | 17-0242-82 |
| CD45 monoclonal (30-F11), PE | eBioscience/ ThermoFisher Scientific | 12-0451-82 |
| CD140a (PDGF Receptor-α) APC | eBioscience/ ThermoFisher Scientific | 17-1401-81 |
| **Mouse reactive antibodies used for magnetic separation** | | |
| TER-119 monoclonal, Biotin | eBioscience/ ThermoFisher Scientific | 13-5921-81 |
| CD16/32, clone 2.4G2, Biotin | BD Biosciences | 553143 |
| CD104 [346-11A] Biotin | BioLegend | 123603 |
| CD31 [MEC13.3] Biotin | BioLegend | 102503 |
| CD45, Biotin | eBioscience/ ThermoFisher Scientific | 13-0451-82 |
| **Other mouse reactive antibodies** | | |
| Ki-67/MKI67 polyclonal | Novus Biologicals | NB500-170SS |
| Podoplanin polyclonal | R&D Systems | AF3244 |
| Phospho-Histone H2A.X (Ser139) | Cell Signaling Technology | 2577S |
| Pro-surfactant Protein C (SFTP-C) polyclonal | Millipore Sigma | AB3786 |
| Platelet-derived growth Factor receptor-α | R&D Systems | AF1062-SP |
| CD45 (30-F11) | Novus Biologicals | NB100-77417SS |
| **Secondary detection antibodies, and reagents** | | |
| Goat anti-rabbit IgG secondary antibody-Alexa fluor 594 | ThermoFisher Scientific | R-37117 |

| | | |
|---|---|---|
| Donkey anti-goat IgG secondary antibody-Alexa fluor 594 | ThermoFisher Scientific | A-11058 |
| Donkey anti-Mouse IgG secondary antibody-Alexa Fluor 594 | ThermoFisher Scientific | R37115 |
| Donkey anti-Rabbit IgG Secondary Antibody-Alexa Fluor 488 | ThermoFisher Scientific | R37118 |
| Anti-Biotin MicroBeads UltraPure | Miltenyi Biotec | 130-105-637 |
| **Fluorescent cell staining reagents** | | |
| Fixable Viability Dye eFluor® 450 | eBioscience/ ThermoFisher Scientific | 65-0863-14 |
| Phalloidin-iFluor 594 Reagent - CytoPainter | Abcam | ab176757 |
| Hoechst 33342 trihydrochloride trihydrate | ThermoFisher Scientific | H3570 |
| **Cell culture media and reagents** | | |
| DMEM (Dulbeccos Modification of Eagles Medium)/F12 50\|50 Mix [-] L-glutamine | Corning | 15-090-CV |
| DMEM, [+] 4.5 g/L glucose, sodium pyruvate [-] L-glutamine | Corning | 15-013-CM |
| XF-DMEM media | Corning | 90-113-PB |
| Phosphate Buffered Saline (PBS), pH 7.4 without calcium and magnesium | Cellgro | 21-040-cv |
| Hanks' Balanced Salt Solution (HBSS) with calcium, with magnesium, no phenol red | Hyclone | SH30268.01 |
| HBSS with calcium, without magnesium, no phenol red | ThermoFisher Scientific | MT-21-022-CV |
| HEPES (1M) | ThermoFisher Scientific | 15-630-080 |
| ACK Lysis Buffer | Fisher Scientific | A1049201 |
| Trypsin 0.25% 1X Soln | ThermoFisher Scientific | SH30042.01 |
| L-Glutamine | Cellgro | 25-005-Cl |

| | | |
|---|---|---|
| L-Glutamine | Gibco | 25030081 |
| Sodium Pyruvate | Millipore Sigma | P8574-100G |
| Fungizone | Gibco | 15290-018 |
| Chemicals, peptides, and recombinant proteins | | |
| Bovine Pituitary Extract | Millipore Sigma | P1476-2.5ML |
| Collagenase, type 4 | Worthington Biochemical | LS004188 |
| Dispase II | Millipore Sigma | 4942078001 |
| Matrigel Membrane Matrix GFR | Corning | 356231 |
| Insulin-Transferrin-Selenium-Ethanolamine (ITS -X) (100X) | ThermoFisher Scientific | 51500056 |
| Human Insulin Soln. | Millipore Sigma | I9278-5ML |
| EGF Recombinant Mouse Protein | ThermoFisher Scientific | PMG8041 |
| STEMRD Y27632 ROCK Inhibitor | ThermoFisher Scientific | 50-175-997 |
| All-trans Retinoic Acid | Millipore Sigma | R2625-100MG |
| Cholera toxin | Millipore Sigma | C8052-5MG |
| Recombinant human TGF-beta1 | PeproTech | 100-21-2ug |
| Horseradish peroxidase | Millipore Sigma | P8375-25KU |
| Homovanillic acid, Fluorimetric reagent | Millipore Sigma | H1252-100MG |
| Hydrogen peroxide solution, 30 % (w/w) | Millipore Sigma | H1009-100ML |
| Oligomycin | Millipore Sigma | O4876-5MG |
| Carbonyl cyanide 4 | Millipore Sigma | C2920-10MG |
| Antimycin A | Millipore Sigma | A8674-25MG |
| 2-Deoxy D-Glucose | Millipore Sigma | D6134-5G |
| **Assay kits used in this study** | | |
| Pierce ™ BCA Protein Assay Kit | ThermoFisher Scientific | PI23225 |
| EZQ™ Protein Quantitation Kit | ThermoFisher Scientific | R33200 |
| MidiMACS Starting Kit (LS) | Miltenyi Biotec | 130-042-301 |
| Senescence Detection Kit | Abcam | Ab65351 |
| Mouse XL Cytokine Array Kit | R&D Systems | ARY028 |
| ADP/ATP Ratio | Abcam | Ab65313 |
| **Mice used in this study** | | |
| C57BL/6J (2 mos) male | Jackson Laboratory | 000664 |

| | | |
|---|---|---|
| C57BL/6J (18 mos) male | National Institute of Aging | N/A |
| Nox4 knockout | Dr. Karl-Heinz Krause, University of Geneva | N/A |
| **Mouse genotyping** | | |
| Transnetyx, Inc. Cordova, TN | | |
| **Anesthesia agent** | | |
| Isoflurane | VetOne® | NDC 13985-528-60 |
| **Equipment** | | |
| Inverted Microscope | Keyence | BZ-X710 |
| Inverted Microscope | Carl Zeiss | Invertoskop 40C |
| Nikon A1R confocal microscope | Nikon | |
| Microplate reader | BioTek | SynergyMx |
| Centrifuge | Eppendorf | 5702R |
| Flow Cell Sorter | BD Biosciences | BD FACS Aria |
| Flow Cell Analysis | BD Biosciences | BD LSR II GUAVA EasyCyte |
| Extracellular Flux Analyzer | Agilent | Seahorse XFe96 |
| **Accessories** | | |
| LS columns | Miltenyi Biotec | 130-042-401 |
| Novex NuPAGE 10% Bis-Tris Protein gels | Invitrogen | NP0315BOX |
| Novex Colloidal Blue Staining kit | Invitrogen | LC6025 |
| Amicon Ultra 4ml, 3kDa molecular weight cut-off filters | Millipore Sigma | UFC800324 |
| Cell culture insert 0.4 um pore size, 24 well | Fisher Scientific | 353095 |
| HistoGel; Specimen Processing Gel | Richard-Allan Scientific/ ThermoFisher Scientific | HG-4000-012 |
| Vectashield | Vector Laboratories | H-1400-10 |
| **Major software used** | | |
| Image J | Wayne Rasband (NIH) | https\|//github.com/imagej/imagej1 |
| Image Quant | Cytiva | *Version* TL8.1 |
| Flow Jo | Tree Star Inc. | |
| Nis Elements | Nikon | *Version* 5.0 |
